# LTMG: A novel statistical modeling of transcriptional expression states in single-cell RNA-Seq data

**DOI:** 10.1101/430009

**Authors:** Changlin Wan, Wennan Chang, Yu Zhang, Fenil Shah, Xiaoyu Lu, Yong Zang, Anru Zhang, Sha Cao, Melissa L. Fishel, Qin Ma, Chi Zhang

## Abstract

A key challenge in modeling single-cell RNA-seq (scRNA-seq) data is to capture the diverse gene expression states regulated by different transcriptional regulatory inputs across single cells, which is further complicated by a large number of observed zero and low expressions. We developed a left truncated mixture Gaussian (LTMG) model that stems from the kinetic relationships between the transcriptional regulatory inputs and metabolism of mRNA and gene expression abundance in a cell. LTMG infers the expression multi-modalities across single cell entities, representing a gene’s diverse expression states; meanwhile the dropouts and low expressions are treated as left truncated, specifically representing an expression state that is under suppression. We demonstrated that LTMG has significantly better goodness of fitting on an extensive number of single-cell data sets, comparing to three other state of the art models. In addition, our systems kinetic approach of handling the low and zero expressions and correctness of the identified multimodality are validated on several independent experimental data sets. Application on data of complex tissues demonstrated the capability of LTMG in extracting varied expression states specific to cell types or cell functions. Based on LTMG, a differential gene expression test and a co-regulation module identification method, namely LTMG-DGE and LTMG-GCR, are further developed. We experimentally validated that LTMG-DGE is equipped with higher sensitivity and specificity in detecting differentially expressed genes, compared with other five popular methods, and that LTMG-GCR is capable to retrieve the gene co-regulation modules corresponding to perturbed transcriptional regulations. A user-friendly R package with all the analysis power is available at https://github.com/zy26/LTMGSCA.

## INTRODUCTION

Single-cell RNA sequencing has gained extensive utilities in many fields, among which, the most important one is to investigate the heterogeneity and/or plasticity of cells within a complex tissue micro-environment and/or development process [1–3]. This has stimulated the design of a variety of methods specifically for single cells: modeling the expression distribution [4–6], differential expression analysis [7–12], cell clustering [13, 14], non-linear embedding based visualization [15, 16] and gene co-expression analysis [14, 17, 18]. etc. Gene expression in a single cell is determined by the activation status of the gene’s transcriptional regulators and the rate of metabolism of the mRNA molecule. In single cells, owing to the dynamic transcriptional regulatory signals, the observed expressions could span a wider spectrum, and exhibit a more distinct cellular modalities, compared with those observed on bulk cells[14]. In addition, the limited experimental resolution often results in a large number of expression values under detected, i.e. zero or lowly observed expressions, which are generally noted as “dropout” events. How to decipher the gene expression multimodality hidden among the cells, and unravel them from the highly noisy background, forms a key challenge in accurate modeling and analyses of scRNA-seq data.

Clearly, all the analysis techniques for single cells RNA-Seq data including differential expression, clustering, dimension reduction, and co-expression, heavily depend on an accurate characterization of the single cell expression distribution. Currently, multiple statistical distributions have been used to model scRNA-Seq data [4, 5, 9, 10]. All the formulations consider a fixed distribution for zero or low expressions disregarding the dynamics of mRNA metabolism, and only the mean expression and proportion of the rest is maintained as target of interest. These methods warrant further considerations: (1) the diversity of transcriptional regulatory states among cells, as shown by the single molecular in situ hybridization (smFISH) data [19–21], would be wiped off with a simple mean statistics derived from non-zero expression values; (2) some of the observed non-zero expressions could be a result of mRNA incompletely degraded, rather than expressions under certain active regulatory input, thus they should not be accounted as true expressions; (3) zero-inflated unimodal model has an over-simplified assumption for mRNA dynamics, particularly, the error distribution of the zero or low expressions are caused by different reasons, negligence of this may eventually lead to a biased inference for the multi-modality encoded by the expressions on the higher end.

To account for the dynamics of mRNA metabolism, transcriptional regulatory states as well as technology bias contributing to single cell expressions, we developed a novel left truncated mixture Gaussian (LTMG) distribution that can effectively address the challenges above, from a systems biology point of view. The multiple left truncated Gaussian distributions correspond to heterogeneous gene expression states among cells, as an approximation of the gene’s varied transcriptional regulation states. Truncation on the left of Gaussian distribution was introduced to specifically handle observed zero and low expressions in scRNA-seq data, caused by true zero expressions, “dropout” events and low expressions resulted from incompletely metabolized mRNAs, respectively. Specifically, LTMG models the normalized expression profile (log RPKM, CPM, or TPM) of a single gene across cells as a mixture Gaussian distribution with K peaks corresponding to suppressed expression (SE) state and active expression (AE) state(s) of the gene. We introduced a latent cutoff to represent the lowest expression level that can be reliably detected under the current experimental resolution. Any observed expression values below the experimental resolution are modeled as left censored data in fitting the mixture Gaussian model. For each gene, LTMG conveniently assigns each single cell to one expression state by reducing the amount of discretization error to a level considered negligible, while the signal-to-noise ratio and the interpretability of the expression data is largely improved. Based on the LTMG model, a differential expression test, a co-regulation module detection and a clustering algorithm were further developed.

A systematic method validation was conducted with the following key results: (1)LTMG achieves the best goodness of fitting in 23 high quality data sets, compared with four commonly utilized multimodal models of scRNA-seq data; (2) using a set of mRNA kinetic data, we confirmed the validity of treating a significant portion of the low but non-zero expressions as a result of un-fully degraded mRNA in LTMG, which should not be considered as true expressions under active regulations; (3) on a cancer single cell RNA-seq data, we demonstrated that single cell groups defined by distinct gene expression states captured by LTMG, are in good agreement with known sub cell types, i.e., exhausted CD8+T cell population and subclasses of fibroblast cells, in other words, the multi-modality setting in LTMG uncovers the heterogeneity among single cells; (4) non-linear embedding and cell clustering based on LTMG discretized expression states produces more informative clusters; (5) we generated a single cell RNA-seq data with perturbed transcriptional regulation and validated the high sensitivity and specificity of the LTMG based differential gene expression and gene co-regulation analysis. A user-friendly R package with all the key features of LTMG model was released through https://github.com/zy26/LTMGSCA.

## METHODS

### Mathematical model linking gene expression states in single cells to transcriptional regulation

A gene’s expression in a mammalian cell is the result of the interactions between its DNA template and a collection of transcriptional regulatory inputs (TRIs) including: (1) transcriptional regulatory factors (TFs) (cis-regulation); (2) miRNA or lncRNA; (3) enhancer and super-enhancer; and (4) epigenetic regulatory signals[22, 23]. For a gene with *P* possible transcriptional regulation inputs, *TRI*_*i*_, *i*= 1, …, *P*, the probability of its promoter being bound by an RNA polymerase, P_b_, which is proportional to the rate of its transcription, can be modeled by a Michaelis Menten equation [24, 25]

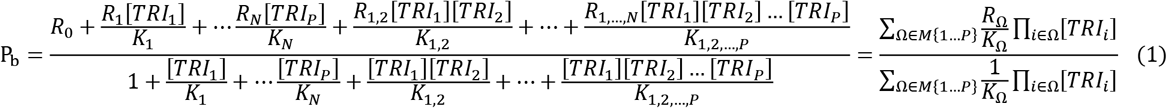

where *R*_*i*_, [*TRI*_*i*_], *K*_*i*_ denote production rate, concentration and kinetic parameters associated with the *i*th TRI; *M*,{1 … *P*} is the power set of {1 … *P*}, *R*_Ω_, *K*_Ω_ denote the production rate and kinetic parameters associated with the interactive effects of TRIs in Ω, where Ω ∈ *M*{1 … *P*}. The set of active TRIs in a single cell fully determines the transcription rate of the gene, and thus its transcriptional regulatory state (TRS). Note that in a single cell each TRI can be rationally simplified to have two states: bound or not bound to the DNA molecule, thus the *TRI*_*i*_ is a Boolean variable and equation (1) becomes a discrete function with at most |*M*{1 … *P*}| = 2^P^ plateau levels:

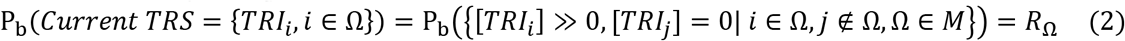

Such discretization of gene’s transcriptional rate greatly simplified the kinetic model and has achieved satisfactory performances in deriving the transcriptional regulatory dependency between the gene’s expression state and its TRIs [26, 27].

For a mammalian cell, the total number of combinations of TRIs can be substantially large, especially considering the epi-genetic regulators[22]. However, the number of TRSs of a gene in a single cell RNA-seq experiment is always much smaller. The reason being: 1) the phenotypic diversity of the cells measured in one experiment is relatively small; 2) local interactive effects among multiple TRIs are exerted on the same regulatory element [23]; and 3) some master repressors such as chromatin folding or certain TFs can dominate the regulation of the gene’s expression[23].

Denote *M*^*X*^ as the set of all possible TRS of gene X and 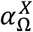 as the probability of sampling a cell with TRS Ω, Ω ∈ *M*^*X*^, from the cell population in a single cell experiment. With introducing a Gaussian error to the discretized model of the formula (2), the probability density function of the transcriptional rate of X in a single cell can be modeled as a mixture Gaussian distribution:

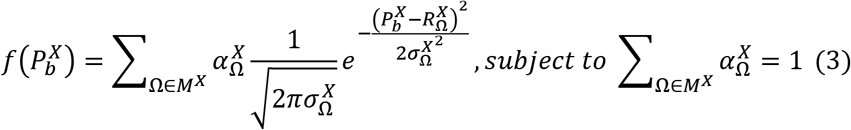

where the mixing probability, mean and standard deviation, 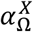, 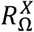 and 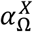 are unknown. Single cell RNA-seq measures the abundance of mature mRNA in cytosol, determined by the transcription and degradation rate of the mRNA. The gene expression pattern we eventually observed is mainly shaped by the (i) cytosol mRNA abundance, compounded by (ii) observation errors and (iii) experimental resolution. Under the assumption of several common transcriptional regulation models, including constant transcriptional regulatory input and transcriptional burst [28], we derived that the multimodality of transcription inputs and rates defined in (2) and (3) can be extended into the multimodality of mRNA abundance with assuming Gaussian observation errors (See more details in **Supplementary Methods**).

Denote 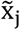, j = 1 … N as the normalized gene expression level (such as log CPM or TPM) of gene X in a scRNA-seq experiment with individual library constructed for N cells and measured with high sequencing depth. Based on the derivations above, we illustrated the relationship between the repertoire of the TRSs of X and its observed gene expression profile in Figure 1A. A mixture Gaussian model is utilized to characterize the distribution of observed normalized gene expression level of X through multiple cells. Gene expressions falling into a same peak are considered to have the same Gene Expression State, that share the same TRS or different TRS with a similar mean pattern; while the expressions falling into different peaks are more likely to have different TRSs. We index the Gaussian peaks by their means and denote the one with smallest mean as the peak 1, and define 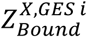 as the boundary between the (*i* + 1)*th* and *ith* peak, which can be estimated by maximizing the likelihood function.

**Figure 1.**
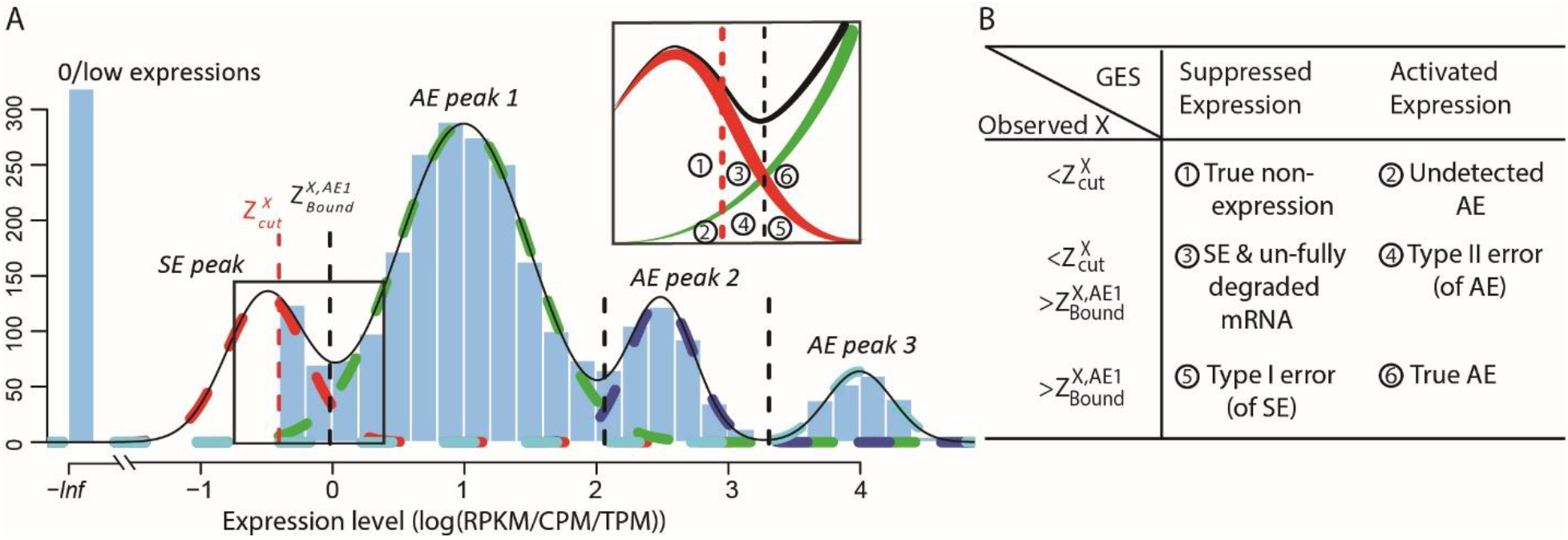
(A-B) The relationship between observed genes expression level, the gene’s SR and AR TRSs, and the experiment resolution threshold 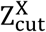. The histogram in light blue illustrates the distribution of the log normalized gene expression (RPKM, CPM or TPM) of one gene in a scRNA-seq data. The four dash curves represent the four fitted mixture components, corresponding to one SE and three AE peaks. 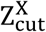 is shown as the red dash line. The framed panel on top right is a zooming in of the non-zero low expression distribution, which is divided into six small areas (B): ① True non-expression, ② expression under suppressed expression & incompletely degraded mRNA, ③ Type I error of SE & incompletely degraded mRNA, ④ Undetected true expression, ⑤ true non-expression but detected as zero and ⑥ True active expression, with detailed definition given in Supplementary Note.

For a robust estimation of the multimodality of the observed expression profile, a key challenge is to address the observed low but non-zero expressions. These low observations could be a result of multiple factors, such as technique errors, un-fully degraded mRNAs and varied experimental resolutions. We introduced a latent threshold 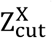 where, when 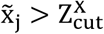, x̃_j_ is modeled by mixture Gaussian distribution; while when 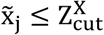, it cannot be reliably quantified under the current experimental resolution. Correspondingly, peaks of mean smaller or larger than 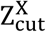 are called suppressed expression (SE) or active expression (AE) peaks. 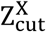 differentiates the large expression values that are more likely to be under active expression state, and those low expression values that are not reliably quantifiable. In scRNA-seq data, other than a small number of housekeeping genes, an SE peak generally exists in the expression profile of most genes.

Figure 1A and 1B illustrates the relationship between the expression states of X, observed expression level 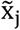, and 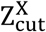. Specifically, when 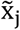 is observed to be zero, it can be ① true non-expression or undetected expressions under an suppressed expression state and ② undetected active expression, i.e. the commonly defined “drop-outs”; when 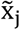 is low but non-zero, its observation can be caused by several reasons including: ③ true zero expression but with a sequencing error, or X is under a suppressed expression in the cell j, and there is incompletely degraded mRNA after the switch from an active expression to the current suppressed expression state and ④ type II error of an active expression state; when 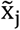 is large, ⑤ 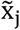 is observed as an type I error of suppressed expression state and ⑥ it is with a high probability that the observed 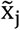 can reflect the true gene’s expression state.

Based on the derivations above, we could model a single cell’s gene expression profile as a multimodal distribution, with observations smaller than 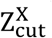 left truncated. Under the current model, active expression state, i.e., the AE peaks, can be robustly inferred; and the unquantifiable the non-zero low expressions, i.e., the SE peak(s), can be effectively handled.

### Left Truncated Mixture Gaussian (LTMG) distribution for gene expression modeling

To accurately model the gene expression profile of scRNA-seq data, we developed a **L**eft **T**runcated **M**ixture **G**aussian model, namely LTMG, to fit the log transformed normalized gene expression measures of gene X, such as TPM, CPM or RPKM, over N cells as X = (*x*_1_, *x*_2_, …, *x*_*N*_). We assume that *x*_*i*_ follows a mixture Gaussian distribution with K Gaussian peaks corresponding to different SE and AE peaks.We introduce a parameter 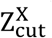 and consider the log transformed zero and low expression values smaller than Z_cut_ as left censored data. With the left truncation assumption, X is divided into reliably measured expressions 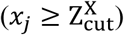 and left-censored gene expressions 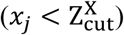. The density function of X can be written as:

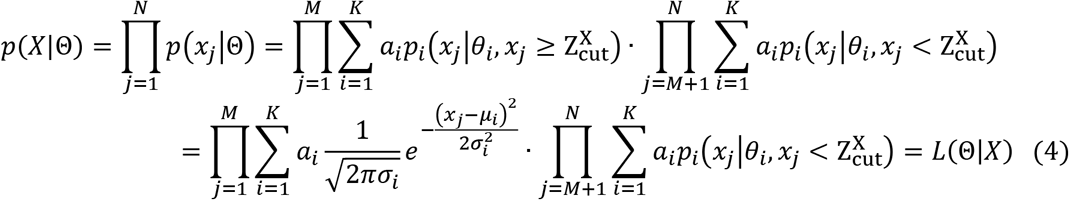

where parameters Θ = {*a*_*i*_, *u*_*i*_ σ_*i*_| *i* = 1 … *K*} and *a*_*i*_, *u*_*i*_ *and* σ_*i*_ are the mixing probability, mean and standard deviation of the K Gaussian distributions, corresponding to K expression states, M is the number of observations *x*_*j*_ that are larger than 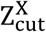, N is the total number of observations. Θ can be estimated using EM algorithm with given 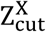 and K. The computation of 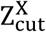 for each gene, EM algorithm for estimating Θ, selection of K, and complete algorithm and mathematical derivations are detailed in **Supplementary Methods**.

### Datasets used for model comparison

To conduct a comprehensive evaluation our model, we collected 23 datasets totaling 66,780 human and mouse cells across different cell extraction and sequencing platforms and with varied experimental designs. It is noteworthy there are multiple scRNA-seq protocols varied by cell capture, lysis and sequencing, which majorly falls into two categories namely individual library for each cell and drop-seq based methods. Recent reviews suggested that the Smart-Seq2 protocols achieve best performance among the methods of the first type while 10x Genomics Chromium is the most utilized commercialized pipeline [3]. Our data collection comprehensively covers human and mouse data generated by Smart-seq/Smart-Seq2, 10x Genomics and Drop-seq platforms from January 2016 to June 2018 on the GEO database. Hence, we consider this collection form an unbiased testing set can represent the general characteristics of the single cell data generated from the two types of protocol. The detailed data information was listed in the supplementary table 1. Since each dataset has different levels of complexity, in order to evaluate the model performances, we generated sub datasets within each of the 23 datasets, so that sub-datasets will have comparable levels of complexities. The sub datasets were extracted to represent three different types of sample complexities: (1) pure condition, where each sub dataset contains cells of one type under a specific experimental condition; (2) cell cluster, where each sub dataset belongs to priori computationally clustered cells; and (3) complete data, where each sub dataset contains multiple mixed cell population, such as cells from one cancer tumor tissue (see detail in Supplementary Methods). In total, 51 pure condition, 49 cell cluster, and 78 complete data sub data sets were extracted from the 23 large data sets. It is noteworthy that all the extracted sub data set are only composed by cells from one of the 23 original data sets. Hence no extra error caused by the batch effect among the data sets needs to be addressed.

### Comparisons of the goodness of fitting of LTMG with ZIMG, MAST and BPSC models

We applied Zero-inflated mixed Gaussian (ZIMG), Left Truncated Mixed Gaussian (LTMG), MAST[4] and Beta Poisson (BPSC)[5] on each dataset, each of which has the following parameter setting. We use MAST with default parameters, and for each gene only non-zero values were used and fitted with Gaussian distribution. For BPSC, to achieve a reliable estimation, only genes with non-zero expressions in at least 25 single cells were kept. ZIMG was used with default parameters. Kolmogorov Statistic (KS) is used to measure gene-wise goodness of fitting. For each gene, the KS score is assessed by using the none zero observations for ZIMG, MAST and BPSC models and normalized by dividing the KS score by the none zero proportions, due to their zero inflation assumption. Only genes kept for all four models are used for downstream evaluations.

For each extracted sub dataset, we defined a goodness fitting score for each method using the mean and standard deviation of gene-wise KS values:

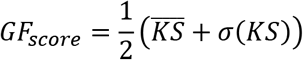

where 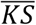 is the mean value of gene-wise KS scores from a dataset and σ(*KS*) the standard deviation. The GF score evaluates each method on both overall accuracy (lower 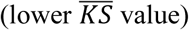 value) and stability (lower σ(*KS*)), and smaller GF indicates better goodness of fitting. The mean and variance of gene-wise KS values for each sub dataset corresponding to all four were all provided in the supplementary table 2.

### Modeling of mRNA metabolic rate with the LTMG model

We collected experimentally measured kinetics of mouse fibroblast cells, particularly the mRNA half-life, of 5028 mRNAs from Schwanhäusser et al’s work [29] and two mouse fibroblast scRNA-Seq datasets [30–33] (GSE99235 and GSE98816). To the best of our knowledge, this is the only cell type with both whole genome level kinetics of mRNA metabolism and scRNA-seq data available in the public domain. In order to pick out the fibroblast cells, we first performed cell clustering using Seurat[34] with default parameters, and each cluster was further annotated with the expression level of fibroblast cell gene markers[35]. In total, we identified 397 fibroblast cells in the GSE99235 and 1100 fibroblast-like cells in GSE98816 datasets. Heatmaps of marker gene expression and t-SNE clustering plots for three datasets were displayed in Supplementary figure 1.

If the hypothesis of the LTMG model is correct, the ratio of the observed low expression caused by un-fully degraded mRNA in the SE peak, which is modeled as 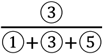 in Figure 1, should be positively correlated with the mRNA half-life, i.e. there is a higher probability to observed low but non-zero expressions for the genes with longer half-life. By applying LTMG on the fibroblast cells extracted from each data set, we tested this hypothesis by measuring the correlation between the mRNA half-life and proportion of uncensored expression in SE peak, i.e. 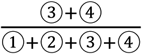, an approximation of 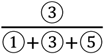. To normalize the impact of the parts ②, ④, and ⑤, i.e. different rate of the type I error of SE peak and the type II error of AS peak of each gene, we compute the correlation conditional to the mean of the first AE peak. Specifically, for each dataset, we ordered genes based on the mean values of their first AE peaks from low to high and split every 100 genes into a group, which gave us 21 and 18 groups in GSE99235 and GSE98816 data sets, respectively. Within each group, Spearman correlation between the mRNA half-life and proportion of uncensored expressions in the SE peak of genes is calculated, and the significance was assessed by using the Student’s T distribution based test.

### Analysis of cell type specifically expressed genes

For any gene, and cells with a priori known cell type identities, since a cell is designated to a peak with largest probability, the peak enrichment score of a cell type is then defined as the exponential function value of the proportion of each cell type falling within a peak type, either SE or AE. The enrichment score is calculated for all cell type gene markers, and due to the specificity of these gene markers, a cell type should have a high AE peak enrichment score for a gene if it is indeed its gene markers, while a high SE peak enrichment score if it is the gene markers for another cell type. The enrichment score is used to evaluate how LTMG model is specific in identifying truly expressed genes.

### T-SNE visualization of the head and neck cancer

We clustered GSE103322[36] datasets by using the Rtsne package with 30 complexity and 20000 max iterations. We only used the markers genes provided by the original paper for cell clustering. The t-SNE analysis is only for data visualization. Cell type annotated in the original work was used to label the cell types.

### LTMG based dimension reduction, visualization, and comparisons with other methods

We applied five dimension reduction methods namely LTMG UMAP, LTMG t-SNE, UMAP, t-SNE and SIMLR on three datasets: GSE103322, GSE72056 and 10x PBMC data set with known cell labels. The LTMG UMAP and LTMG t-SNE methods were conducted with LTMG inferred gene expression states as the input, by using R UMAP package with the default parameters and RTSNE function with perplexity=30 and max iteration=20000; the UMAP and t-SNE methods used original expression data as input (CPM/RPKM) with the same parameters; and the SIMLR method used original expression data as input with default parameters [16]. For the LTMG based inference of expression states, we first compute the SE or AE peaks of a gene’s expression profile and assigning its expression state in each cell by the index of the peak that its expression value with the maximal likelihood. Specifically, an expression value is discretized as an integer k if it is most likely to be assigned to the kth AE peak (k>0) or the SE peak (k=0). When applying SIMLR, we first determined the cluster number ranged from 5 to 15 by using the SIMLR built-in function SIMLR_Estimate_Number_of_Clusters. The number was further used in the clustering analysis of SIMLR.

We evaluated the clustering performance by sum of silhouette width of all the cell (See details in Supplementary Methods). Cell type information are directly retrieved from original works or related sources. Since GSE103322 and GSE72056 provides a comprehensive list of cell marker genes, we conducted dimension reduction and cell clustering by using the marker genes.

### LTMG based differential expression analysis

Under the framework of LTMG, we define that a gene is differentially expressed between the cells of two conditions, if at least one gene expression state (either SE or AE) of the gene has a significantly different representing level in one condition versus the other. Our comprehensive analysis revealed that on average more than 83.8% genes in the PC and CC groups are fitted with one and two peaks, which can be well fitted by a LTMG-2LR model with a modified EM algorithm (**Supplementary Note**). We perform DGE differently for genes either fitted with LTMG-2LR distribution or not, on samples pooled over all conditions. For a given gene X in a scRNA-seq data under J conditions, denote 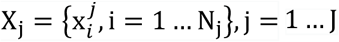 as its expression profile in the N_j_ cells of the jth conditions. The following pseudo codes illustrate our differential gene expression analysis approach, namely LTMG-DGE.

If the gene is fitted with LTMG-2LR distribution. In this case, we assume a gene shares the same SE state and similar degradation rates through different conditions. And we test the differences in proportion and mean of the AE peaks of different conditions. For X_j_, j = 1 … J, we first fit an LTMG-2LR for each X_j_ assuming the same 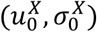 of the SE peak through all the conditions, namely:

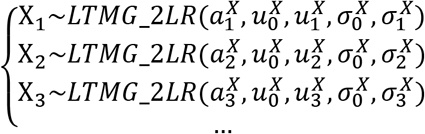

Then differences in 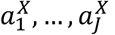 and 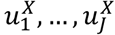 can be rigorously tested by implementing a GLM model with a random sampling process as detailed below. With 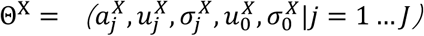 estimated, the probability that 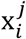 belongs to a SE (or AE) peak can be assessed, denoted as 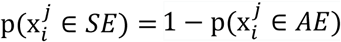. A sampling process can be made by randomly assigning 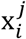 to the SE (or AE) state of condition j with probability 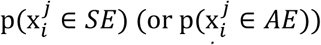, by which 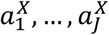 can be tested by using a logit linking function to link the frequency of 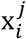 belong to the SE (or AE) state under each condition, with the design matrix of the conditions; and 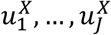 can be tested by using a linear linking function to link the mean of 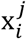 belong to the AE state under each condition, with the design matrix. Applying the random sampling process N times, p value of each test is estimated by the median of the identified p values, and the confidence intervals of each p value can be estimated. The advantages of this process include (1) rigorousness of the GLM form, (2) high sensitivity for the changes in frequency or mean expression level of the AE peak, and (3) the testing rigorousness is not affected by the dilemma of a mixture distribution, due to 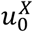 and 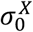 are fixed for all conditions.

If the gene is fitted with more than two AE peaks in at least one condition. We applied the following hypergeometric test based DGE test: (1) fit an LTMG model by using the data of all conditions, i.e. X~*LTMG*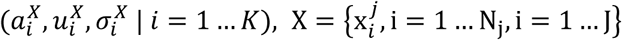, (2) compute the likelihood that 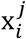 belongs to peak *i* = 1 … *K* and assign 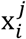 to the peak with the maximal likelihood, (3) compute if the samples of each condition *j* = 1 … *J* are enriched to a peak *i* = 1 … *K* via a hypergeometric test.

The difference of the two testing schemes is that the former one assumes a gene has only one AE peak in each condition, which can vary in proportion, mean, or variance through different conditions, and the test is done on the proportion and mean of the AE peak, while the later fits one LTMG model over the pooled data through all conditions, and test if one condition is specifically enriched with one expression state.

### Single Cell RNA-Sequencing

Pa03C cells were obtained from Dr. Anirban Maitra’s lab at The Johns Hopkins University[37]. All cells were maintained at 37°C in 5% CO2 and grown in DMEM (Invitrogen; Carlsbad, CA) with 10% Serum (Hyclone; Logan, UT). Cell line identity was confirmed by DNA fingerprint analysis (IDEXX BioResearch, Columbia, MO) for species and baseline short-tandem repeat analysis testing in February 2017. All cell lines were 100% human and a nine-marker short tandem repeat analysis is on file. They were also confirmed to be mycoplasma free.

Cells were transfected with either Scrambled (SCR) (5′ CCAUGAGGUCAGCAUGGUCUG 3′, 5′ GACCAUGCUGACCUCAUGGAA 3 ′) or siAPE1 (5 ′ GUCUGGUACGACUGGAGUACC 3 ′, 5 ′ UACUCCAGUCGUACCAGACCU 3′ siRNA). Briefly, 1×10 ^5^ cells are plated per well of a 6-well plate and allowed to attach overnight. The next day, Lipofectamine RNAiMAX reagent (Invitrogen, Carlsbad, CA) was used to transfect in the APE1 and SCR siRNA at 20 nM following the manufacturer’s indicated protocol. Opti-MEM, siRNA, and Lipofectamine was left on the cells for 16 h and then regular DMEM media with 10% Serum was added.

Three days post-transfection, SCR/siAPE1 cells were collected and loaded into 96-well microfluidic C1 Fluidigm array (Fluidigm, South San Francisco, CA, USA). All chambers were visually assessed and any chamber containing dead or multiple cells was excluded. The SMARTer system (Clontech, Mountain View, CA) was used to generate cDNA from captured single cells. The dscDNA quantity and quality was assessed using an Agilent Bioanalyzer (Agilent Technologies, Santa Clara, CA, USA) with the High Sensitivity DNA Chip. The Purdue Genomics Facility prepared libraries using a Nextera kit (Illumina, San Diego, CA). Unstrained 2×100 bp reads were sequenced using the HiSeq2500 on rapid run mode in 1 lane.

### qRT-PCR

qRT-PCR was used to measure the mRNA expression levels of the various genes identified from the scRNA-seq analysis. Following transfection, total RNA was extracted from cells using the Qiagen RNeasy Mini kit (Qiagen, Valencia, CA) according to the manufacturer’s instructions. First-strand cDNA was obtained from RNA using random hexamers and MultiScribe reverse transcriptase (Applied Biosystems, Foster City, CA). Quantitative PCR was performed using SYBR Green Real Time PCR master mix (Applied Biosystems, Foster City, CA) in a CFX96 Real Time detection system (Bio-Rad, Hercules, CA). The relative quantitative mRNA level was determined using the comparative Ct method using ribosomal protein L6 (RPL6) as the reference gene. The primers used for qRT-PCR and qRT-PCR experimental data are detailed in Supplement Table 3. Experiments were performed in triplicate for each sample. Statistical analysis performed using the 2−ΔΔCT method and analysis of covariance (ANCOVA) models, as previously published[38].

### LTMG based gene coregulation analysis

By the formulation of LTMG, for a gene with one SE peak and K-1 different AE peaks, its expression profile across different single cells is modeled by a mixture of K Gaussian distributions; and (2) for a group of genes co-regulated by a specific TRS, each gene’s expression profile, in those cells regulated by the TRS, forms a unimodal Gaussian distribution, after involving independent Gaussian errors. Hence a gene co-regulation model corresponds to a submatrix enriched by 1s in the Binary matrix *M* constructed in the following way:

For a gene X’s expression profile through N samples fitted with one SE and K-1 AE peaks, denote, *P_i_^x^* = 0, 1 … *K* − 1, *i* = 1, …, *N* as the peak with highest likelihood

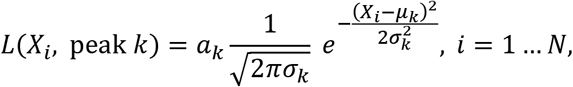

in which 0 represents the SE peak and 1 … *K* − 1 represents the AE peaks. Then a (*K* − 1) × *N* binary matrix 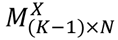 can be constructed by

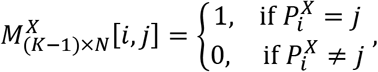

*i* = 1 … *N*, *j* = 1 … *K* − 1. *M* is merged by *M*^*X*^ for the *X* with at least one AE peak. A bi-cluster enriched by 1s in *M* corresponds a group of genes and cells, each of the gene is regulated by one specific TRS through the cells, which is potentially a gene co-regulation module.

We applied our in-house bi-clustering method QUBIC[17, 39] on the binary matrix constructed as above, to identify gene co-regulation modules, namely LTMG-GCR. Specifically, QUBIC is implemented with the following parameters: −o 3000 -f 0.25 -c 0.95. LTMG-GCR is applied to a scRNA-seq data of APEX/Ref-1 KD experiment. Pathway enrichment analysis of the genes in the identified bi-clusters are computed using hypergeometric test against the 1329 canonical pathway and 658 validated transcriptional regulation pathways in MsigDB database[40], with p<0.001 as a significant cutoff.

## RESULTS

### LTMG model substantially improved the goodness of fitting and accurately captured multimodality of scRNA-seq data

To compare the performances of LTMG versus other methods, we collected a comprehensive set of human and mouse scRNA-seq data sets available in the NCBI GEO dataset deposited from January 2016 to June 2018, that were generated by using Smart-seq, Smart-seq2, 10x Genomics and Drop-seq protocols with sample sizes between 100 to 7500 cells. This resulted in 23 data sets totaling 66,780 single cells (Supplementary Table 1). Using the 23 datasets, we further constructed three classes of 178 refined testing data sets, namely (*i*) 51 pure condition datasets, (*ii*) 49 cell cluster datasets and (*iii*) 78 complete data sets, to demonstrate the necessary of a multi-modal assumption in modeling scRNA-seq data. Specifically, the pure condition sets are composed by data of a pure cell type under a tightly controlled condition; cell cluster consists data of computationally predicted cell clusters; and complete data consists data of multiple cell types (supplementary methods).

We first applied LTMG, ZIMG (zero-inflated Mixture Gaussian), MAST (zero-inflated Gaussian), and BPSC (Beta-Poisson) to fit the expression profile of each gene in 91 out of the newly constructed 178 data sets. Kolmogorov Statistics (KS) [41] was applied to evaluate the goodness of fitting of each gene, and for each dataset using each method. Our analysis suggested that LTMG has the significantly better goodness of fitting compared with BPSC and MAST in all the analyzed data sets and outperforming ZIMG in most of the datasets (Figure 2A). We also found that LTMG generally has a smaller number of outliers with poor fitting through all the datasets, (Figure 2B and Supplementary table 4), suggesting that it is more robust comparing to others. Our analysis suggested that the average proportion of genes fitted with one, two, and more than two peaks are 42.5%, 44.9% and 12.6% in pure condition, 16.6%, 65.7% and 17.6% in cell cluster, and 25.4%, 51.5% and 23.1% in complete data sets, respectively.

**Figure 2.**
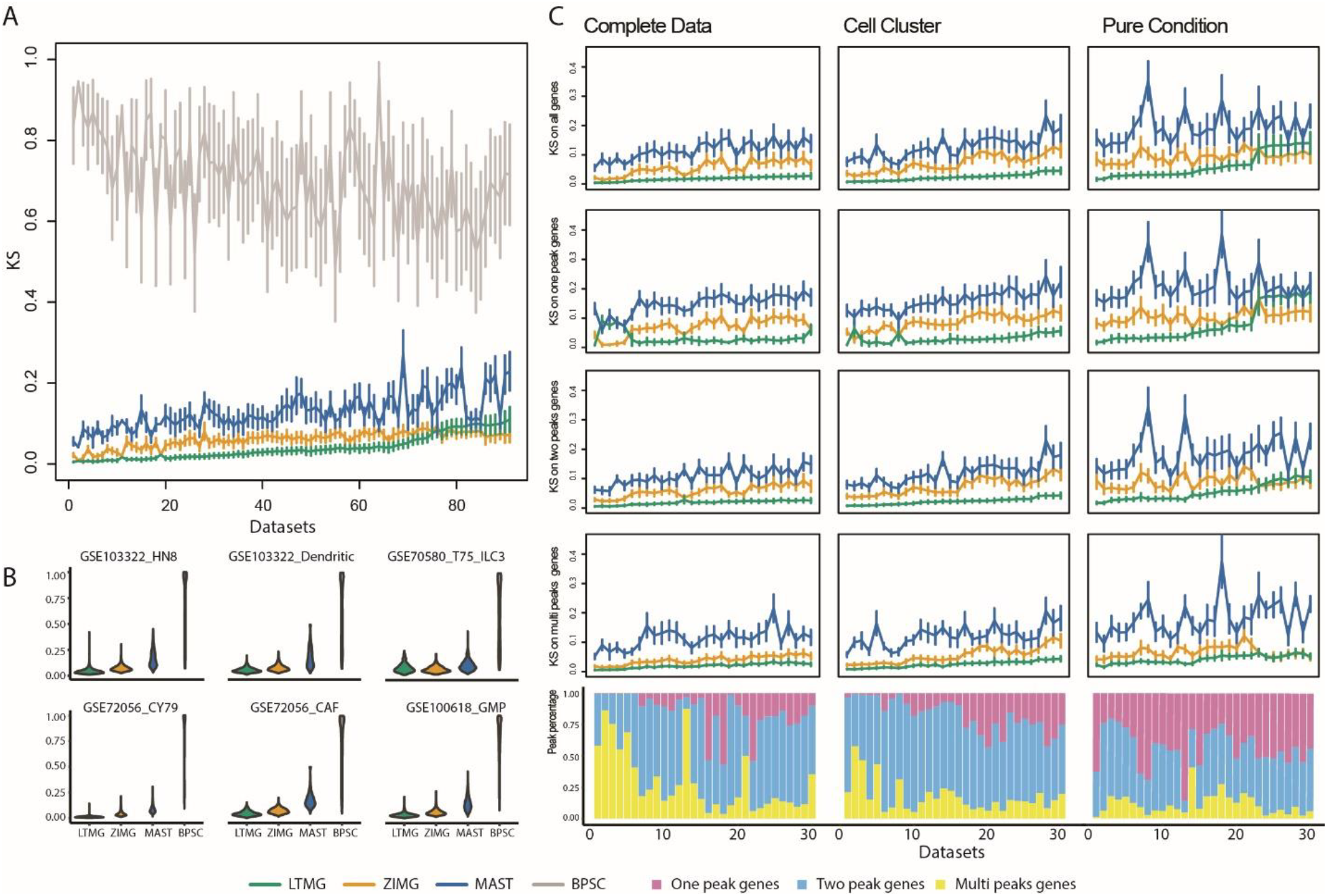
Detailed fitting comparison of LTMG and other models. (A) Goodness of fitting of the four models. X-axis represents different data sets, and Y-axis the goodness of fitting evaluation for each method using KS values, where the mean and standard deviations of the KS values are shown. Note smaller KS values indicate better goodness of fitting. (B) Violin plot of KS value of selected example datasets, 2 for each group. (C) Detailed comparisons of the three models on genes of different peaks and datasets of different groups. The three columns from left to right are the KS values and distribution of peaks in the top 30 complete, cell cluster and pure condition data sets ordered by the KS of LTMG. Horizontal lines in the KS plots represents the mean of KS fitting of value in that group of genes and vertical line is the standard deviation accordingly. Stocked histogram illustrates the percentage distribution of genes of different peaks in different datasets.

Instead of investigating the goodness of fitting over all the genes, we focused on a more detailed comparison of genes that are found to have certain numbers of peaks under LTMG in different groups of testing data. Here we only focused on the comparisons of LTMG with ZIMG and MAST, since BPSC is with much lower performance than these models. In figure 2C, we illustrate the comparison of 30 datasets for each data group, with the performance of rest datasets provided in Supplementary figure 2. Within the cell cluster and complete data sets, LTMG consistently outperformed ZIMG (120/127) and MAST (127/127). In the pure condition datasets, LTMG outperformed MAST in all the tested data sets (51/51), outperformed ZIMG (42/51) for the genes fitted with more than two Gaussian peaks, and have comparable performance as ZIMG (23/51) for the genes that are fitted with one or two peaks. A possible reason for the less significant performance of LTMG on the pure condition datasets could be that the sample size of the PC datasets is generally small (~115 cells on average) compared to cell cluster (~ 388 cells) and complete (~622 cells) data sets. A consequence is that the half bell shaped SE peak (Figure 1A) is not significantly different from a full Gaussian peak when the sample size is small. Notably, ZIMG tends to overestimate the number of AE peaks and overfit the data due to it omits the non-zero expression (falsely) contributed by incompletely degraded mRNA, while LTMG can effectively handle the non-zero low expressions by the left truncation assumption.

To further investigate the models’ robustness in casting the true gene expression levels, we compared the consistency between the pdf of scRNA-seq data inferred by LTMG, ZIMG and MAST models with density of gene expression characterized by matched single cell fluorescence in situ hybridization (scFISH) data. Here we used the only data set with both scRNA-seq and scFISH conducted over the same cell conditions and compared the 15 genes that have been quantified by both of scFISH and scRNAseq. We evaluated the consistency between the pdf of scRNA-seq data and density of scFISH data by using a KL divergence, the higher value of which indicates the better mapping to scFISH data (supplementary methods). Specifically, LTMG achieved larger KL divergence comparing to ZIMG and MAST in 10 genes out of the 15 genes while the three methods achieved similar KL divergence in the rest 5 genes (Supplementary figure 3A). Further visualizations of the expression profile suggested that the multimodality inferred by LTMG is with a higher concordance with the observed expression profile, comparing to other two methods (Supplementary figure 3B).

In addition, we applied the LTMG model to three recent data sets of purified T cells collected from liver, lung and colon cancer tissues [42–44]. These data sets are with large sample size of one purified cell type (5063, 11138, and 12346 cells), hence the distribution of the number of SE and AE peaks derived from these data sets can demonstrate the multi-modality of single gene’s varied expression states in a same cell type. In these data sets, LTMG also achieved the best goodness of fitting. LTMG identified more than 44.5%(4893/10874), 69.73%(7093/10172) and 69.95%(7551/10794) of significantly expressed genes are with at least one SE peak and two AE peaks in the liver, lung, and colon cancer data, respectively (Supplementary figure 4). We further utilized a stringent threshold to identify the genes with at least two AE peaks, each of which covers significant proportion of the total cells and is distinct to other peaks. (see more details in the Supplementary Method). We identified more than 26.56%(2888/10874), 22.67%(2306/10172) and 24.56%(2651/10794) of the significantly expressed genes are with at least two distinct AE peaks in the three data sets, hence further demonstrated the prevalence of observable multiple expression states in large data sets.

### LTMG handles zero and low expressions properly

The observed uncensored low expression depicted as ③ and ④ in Figure 1A are generally seen in all the analyzed data sets, which on average take 27.9%, 16.3% and 14.5% of non-zero values in the PC, CC and CD data (Supplementary Table 5). We hypothesized that one major contributor of the uncensored low expression is the incompletely degraded mRNA under the regulation of a TRS of suppressed state, which should be distinguished from those TRSs under activated states ⑥ (Figure 3A). To validate this hypothesis, we collected a data set of experimentally measured mRNA kinetics of mouse fibroblast cells [29], and two scRNA-seq data set (GSE99235 and GSE98816) of mouse fibroblast cells [31, 45]. We examined the correlations between the mRNA half-life and the estimated proportion of incompletely degraded mRNA. Specifically, positive correlations between (i) the proportions of uncensored observations in the SE peak, defined by 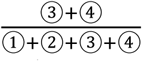 in Figure 1A, and (ii) mRNA half-life, were consistently observed in both data sets (Figure 3B), suggesting that genes with more uncensored expressions regulated by suppressing regulators are probably a result of longer mRNA half-life. It is noteworthy the expression activating peaks with a larger mean may have less impact to the falsely identified non-zero expressions, as the high AE peak illustrated in Figure 3A. To adjust for this bias, we examined the correlations of mRNA half-life with the proportion of uncensored observations with respect to the mean of AE peak (**Methods**). Significant positive correlations (p<0.05) were observed for the genes with a relatively larger mean of AE peak, and the correlations tend to be stronger among the genes with larger AE peaks, in both of the analyzed data sets (Figure 3C), further validated the relationship between the observed low expression and un-fully degraded mRNA.

**Figure 3.**
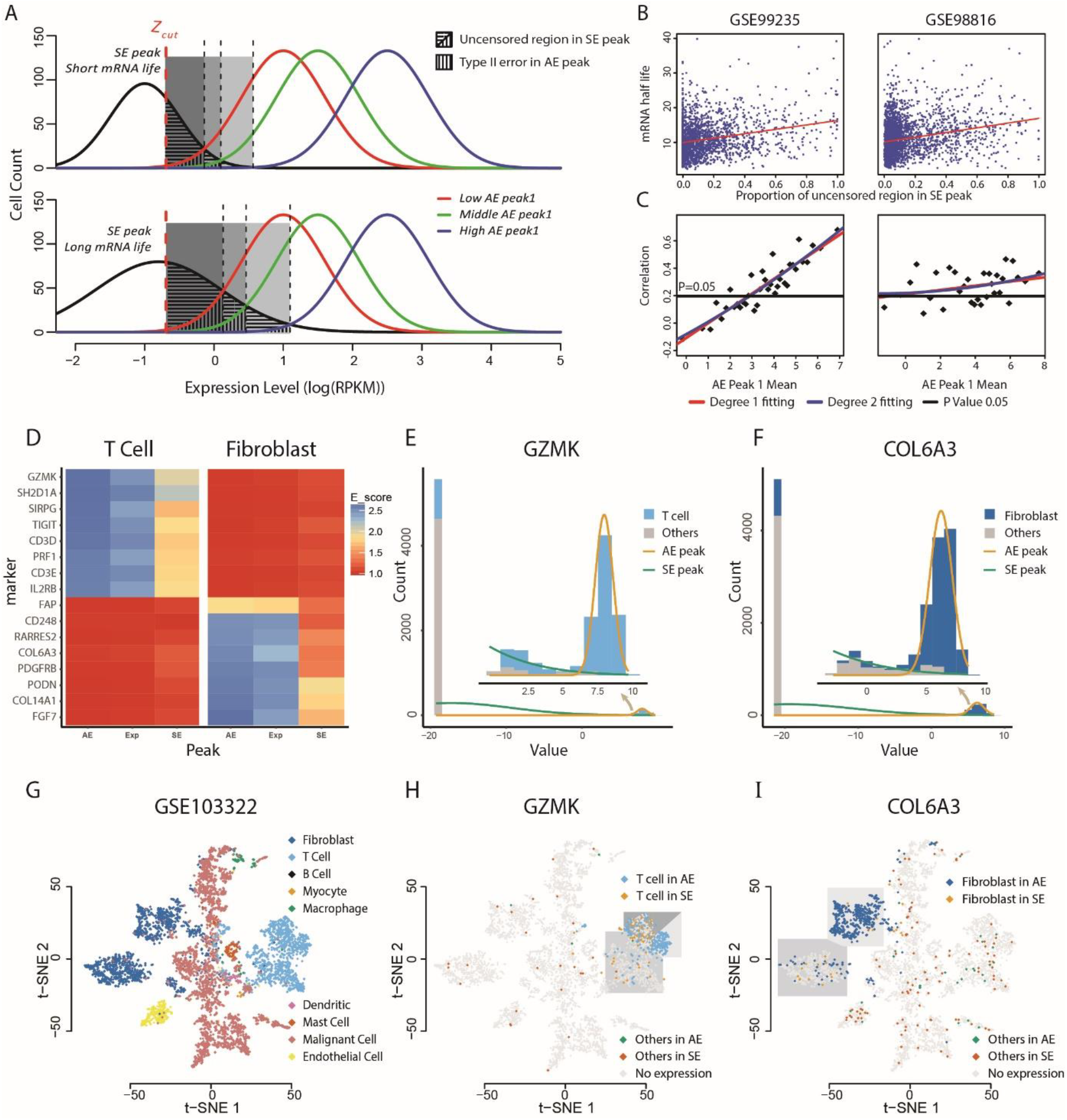
(A-C) Association between the scRNA-Seq measured expression and mRNA degradation rate. (A) Schematic of the uncensored region of genes with different SE peak and influences from different AE peak1. Genes with longer mRNA life tend to have a larger uncensored region. Lower AE peak1 is more likely to introduce a bigger Type II error. (B) Scatter plot of the uncensored region and mRNA half-life in three different datasets. Red line is the degree 1 fitting. (C) Scatter plot of correlation value in different AE peak1 Mean. Red line is degree 1 fitting, blue line is degree 2 fitting, and black line is the correlation threshold when the P value is equal to 0.1. (**D-I) Distribution of AE and uncensored SE expression of cell type markers through different cell types.** (D) Heat map of T cell and fibroblast enrichment information across T cell and fibroblast markers, AE, Exp and SE on the x-axis represents AE peak, non-zero expressions, and non-zero expressions in SE peak. (E, F) Cell distributions with respect to the gene expression and peak fittings of GZMK and COL6A3. Light blue region presents T cells, dark blue presents Fibroblast cells and gray represents other cells. (G) t-SNE plot of different cell types in the GSE103322 dataset. (H) Detailed gene expression states of GZMK in three subclasses of T cells and other cells over the t-SNE plot. (I) Detailed gene expression states of COL6A3 in two subclasses of Fibroblast cells and other cells over the t-SNE plot.

### Modeling the transcriptomic heterogeneity among cells

The multi-modality characteristic of LTMG unravels the transcriptomic heterogeneity through different cells. Next, we ask how cells behave with respect to our identified SE and AE peaks. Our hypothesis is that for the cells with a certain identity such as cytotoxic T cells, they are expected to overly express specific cell marker genes like granzymes, such that their expression level are more likely to be in an AE peak rather than a SE peak in cytotoxic T cells. On the other hand, T cells are more likely to be enriched in certain AE peaks of granzymes but are excluded in SE peaks. In addition, since LTMG identifies certain low non-zero expressions to SE peak, we hypothesize that a cell type will be more strongly enriched to the AE peaks rather than all the cells with non-zero expression value of a marker gene. For a gene, we denoted the cells with non-zero expression of the gene as “Exp”, the cells assigned to the AE peaks as “AE” and the cells assigned to the SE peaks as “SE”. We tested how the cells of different types are distributed through the “AE”, “Exp” and “SE” cell groups of different marker genes.

To conduct the analysis, we applied LTMG on a head and neck cancer (HNSC) data set (GSE103322) consisting 5,902 cells of 9 cell types namely B cell, T cell, Myocyte, Macrophage, Endothelial, Dendritic, and Mast cell, with pre-annotated cell labels and uniquely expressed maker genes identified[1]. We defined an enrichment score to evaluate the association between cell type and the “AE”, “Exp” and “SE” cell groups of each marker gene (methods). Non-surprisingly, our analysis identified that all the cell types are significantly more enriched to the “AE” cell group than the “Exp” and “SE” groups of its marker gene, suggesting that the AE state identified by LTMG better characterizes the true active expressions of the marker genes, comparing to the “only-0” or “fixed-Poisson” consideration of dropout events characterized by MAST or SCDE (Supplementary Table 6). Figure 3D shows the enrichment score of the AE peak, total non-zero expression value and uncensored part in the SE peak of 16 cell markers in the data of T and fibroblast cells. Figure 3E and 3F illustrate the fitted peak distribution of a cytotoxic T cell marker GZMK and a fibroblast marker COL6A3. We further examined the distribution of the AE expression and uncensored SE expression of these two genes in the 2D-tSNE distribution of the cells derived by the complete data (Figure 3G). We observed that the CD8+ T cells with the AE expressions or uncensored SE expressions of GZMK were clearly separated to high cytotoxic and exhausted CD8+ T cells in the HNSC microenvironment[46–48] (Figure 3H). Similarly, the fibroblast cells with an AE or an uncensored SE expression of COL6A3 were differentially distributed as two sub fibroblast types (Figure 3I). Moreover, cells that expressed in SE peak are scattered outside T cell or Fibroblast cell region, validated that SE peak does not representing cell type identity and should be de-noised for further analysis.

### Single cell clustering based on inferred modality by LTMG

Our analysis suggested that the gene expression states inferred by LTMG can reflecting the cell type specific gene expression characteristics with effectively removing the noise of the low but non-zero expressions. Here we show that this denoising approach can promote the dimension reduction based cell clustering analysis and visualization of the single cell data collected from complicated microenvironment such as cancer and peripheral blood samples.

Five dimension reduction and clustering methods including UMAP and t-SNE on the original gene expression data (normalized by TPM/CPM/RPKM) and LTMG denoised data, and SIMLR on original data were compared on three datasets: GSE103322, GSE72056, and 10X PBMC with annotated cell types (**Methods**). We compared LTMG UMAP, LTMG t-SNE, UAMP, t-SNE and SIMLR by using the Silhouette width. the higher value of which suggests a better consistency between predicted cell clusters and true cell labels. Visualization of the 2D embedded data, cell clustering and the silhouette width (sil value) were shown in Figure 4. Our analysis suggested the cell clusters inferred from LTMG denoised data outperform the clusters identified by using original data, for both UMAP and t-SNE based dimension reduction and clustering. In the GSE72056 and GSE103322 dataset, cell surface markers and predicted copy number variations were used to identify true malignant cells, which were composed by multiple subclasses of cells due to inter-tumor heterogeneity, as illustrated by the red colored cells in Figure 4. We observed the malignant cells, as well as other normal cells, are more spreaded over the 2D UAMP and t-SNE of the original data while the LTMG UMAP and LTMG t-SNE well manage the subclass of malignant cells from different patients (Figure 4 and Supplementary Figure 5). In addition, different types of immune and stromal cells were better distinguished from malignant cells and each other in the LTMG UMAP and LTMG t-SNE based embedding. A possible explanation is that the LTMG based transformation of gene expression states can better characterize the inter-cell type varied expression states via removing the intra-cell type gene expression variations that do not form varied expression states.

**Figure 4.**
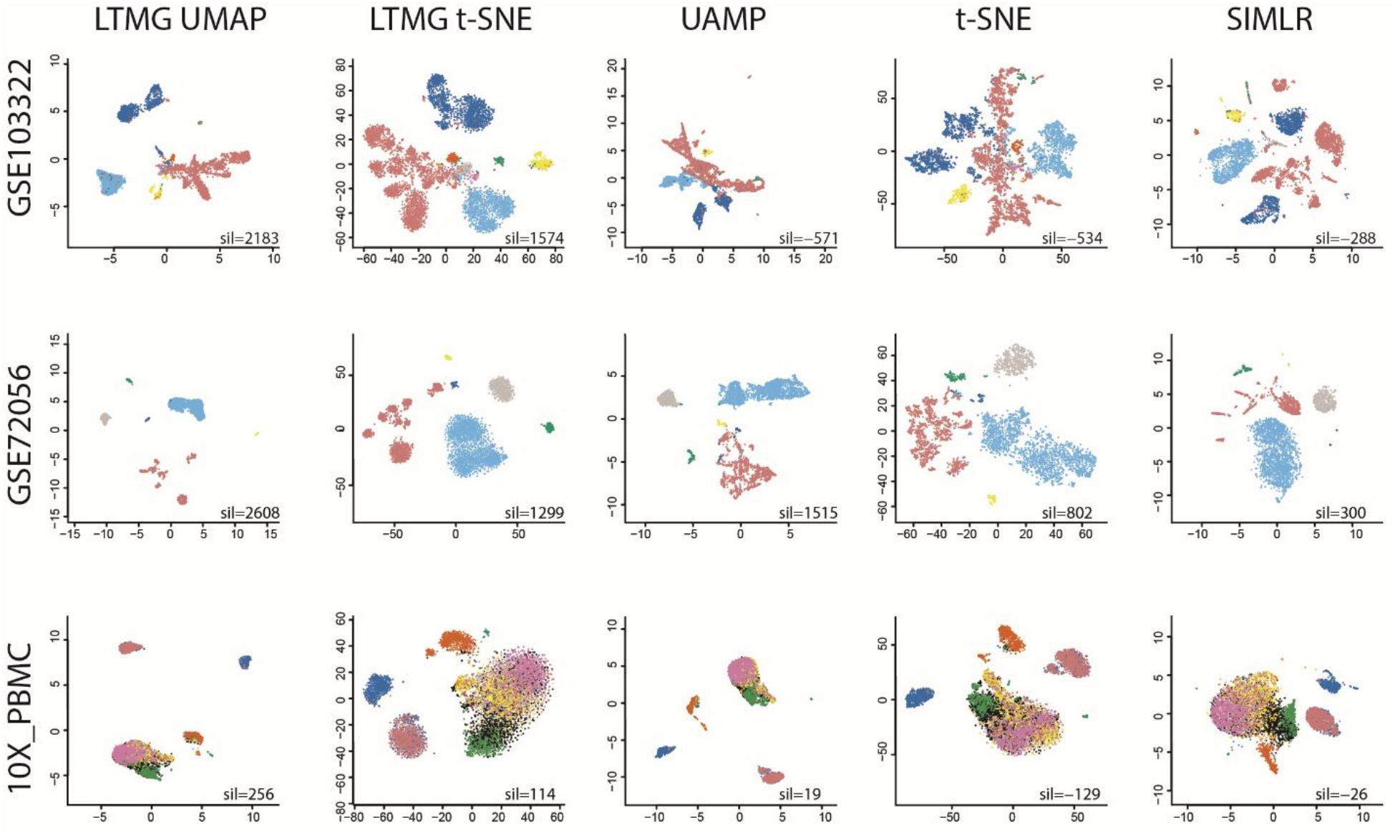
Clustering visualization of three datasets using five methods. 2D visualization of the three datasets GSE103322, GSE72056 and 10X_PBMC embedded by LTMG UMAP, LTMG t-SNE, UMAP, t-SNE and SIMLR. Cells are colored by the cell types annotated in original work. Sil value represent the sum of silhouette width between the predicted cell clusters and known cell labels.

### Differential gene expression and gene co-regulation analysis and experimental validations

Under the formulation of LTMG, a gene is considered as *differentially expressed* between the cells of two conditions if (1) the proportion of the SE or AE peak or the mean of the peak are significantly different between the conditions when both conditions have at most one AE peak, and (2) the proportion of the SE peak or at least one AE peak is significant different between the conditions, when there are more than one AE peaks in one condition (**Methods**). *A gene co-regulation module* can be defined by a group of genes sharing a common TRS throughout a subset of cells. The LTMG based differential gene expression analysis (LTMG-DGE) is further empowered to handle more complicated design by incorporating a generalized linear model setting; and the gene co-regulation analysis (LTMG-GCR) is further equipped by implementing a bi-clustering algorithm to detect co-regulation modules of potential transcriptional heterogeneity[17, 18] (**Methods**). To experimentally validate the LTMG based DGE and GCR analysis, we generated a scRNA-seq data set consisting of 142 patient-derived pancreatic cancer cells under two crossed experimental conditions: APEX1 knockdown (APE1/Ref-1-KD) or control, and under hypoxia or normoxia conditions.

We compared the distribution of differentially expressed genes and their functional relevance to APE1 identified by LTMG-DGE with MAST, SCDE, SC2P, EdgeR and DESeq. Using LTMG-DGE, we identified 448 up- and 1,397 down-regulated genes in APE1/Ref-1-KD vs. control under hypoxia, and 471 up- and 992 down-regulated genes under normoxia (p<0.01); while MAST identified 282 and 521 up-regulated and 397 and 607 down-regulated genes, under hypoxia and normoxia conditions, respectively (p<0.01). In addition, under the hypoxia condition, 215, 187, 129 and 500 up- and 281, 1528, 188 and 1085 down-regulated genes were identified by SCDE, SC2P, EdgeR and DESeq (p<0.01), respectively. The differentially expressed genes identified by the methods are given in Supplementary Table 7. Consistency of the LTMG-DGE and MAST identified differentially expressed genes are shown in Figure 5A and Supplementary Table 7.

**Figure 5.**
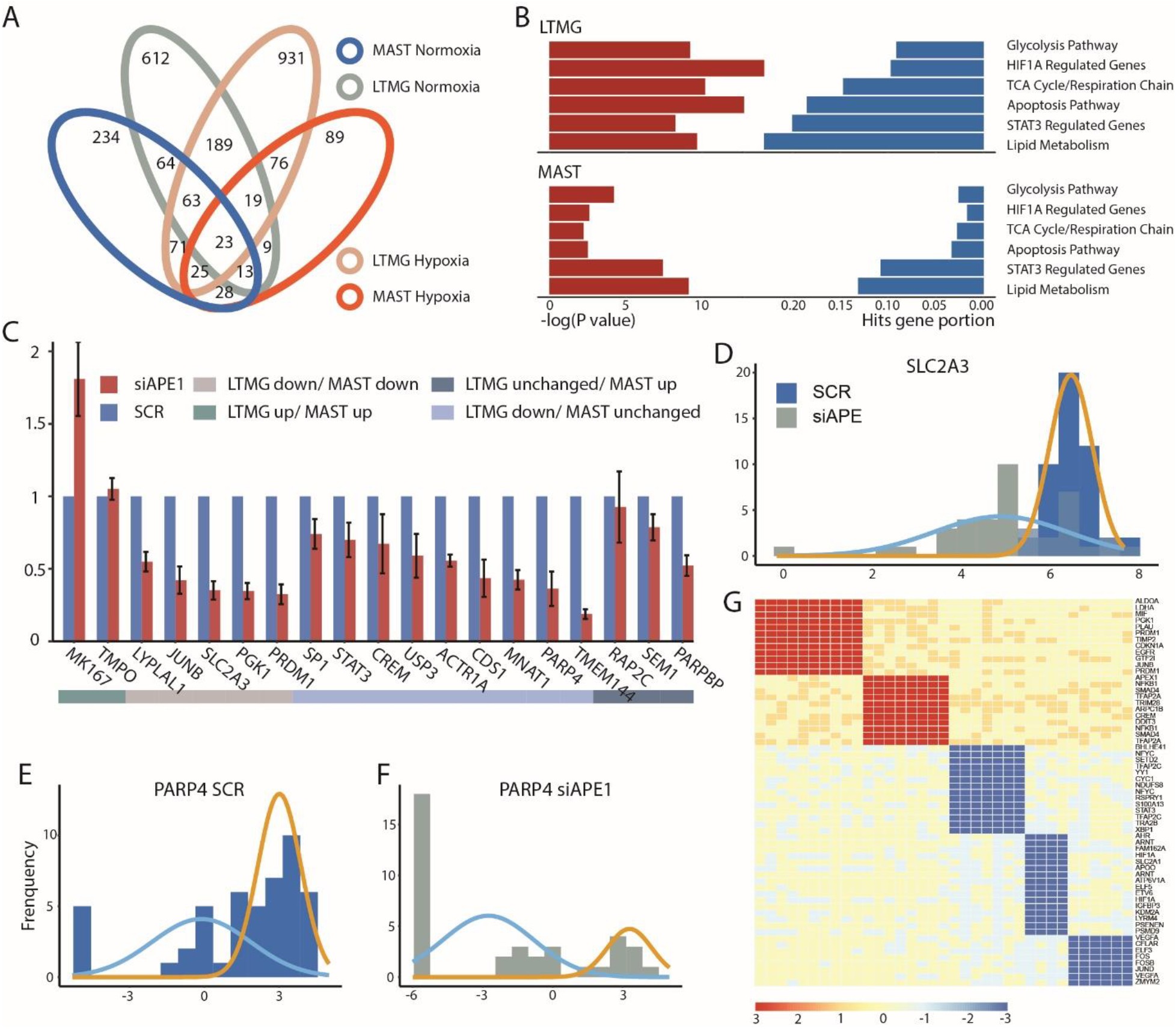
Experimental validation of LTMG-DGE. (A) Overlap of down-regulated genes in APE1/Ref-1-KD vs. SCR control in hypoxia and normoxia, identified by LTMG-DGE and MAST. (B) Enrichment of the genes down-regulated in APE1/Ref-1-KD vs. SCR control in key APE1/Ref-1 related pathway, under hypoxic conditions. (C) Expression of selected genes analyzed by qPCR of Pa03C cells transfected with APE1/Ref-1 siRNA and placed under hypoxia for 24 h. (D-F) Expression profile of SLC2A3 and PARP4 in APE1/Ref-1-KD (siAPE) and control (SCR) under hypoxia. Gene expression level is quantified by log(RPKM) and represented on the x-axis. Gold and blue curves represent peaks correspond to different TRSs. (G) Bi-cluster structures of gene coregulation modules enriched by STAT3 and HIF1A regulated genes. The x-axis represents samples and y axis represents genes. AE and SE status of a gene in a sample are colored by red and blue, respectively.

*APEX1* is a multifunctional protein that interact with multiple transcriptional factors (TFs) to regulate the genes involvement in response to DNA damage, hypoxia and oxidative stress[49]. Our previous study identified significant roles of APEX1 in the regulation of Pa03c cell’s response to microenvironmental stresses[50]. Functional enrichment of the differentially expressed genes identified by the methods were further examined. Comparing to MAST, SCDE, SC2P, EdgeR and DESeq, the down-regulated genes in APE1/Ref-1-KD vs. control under hypoxia conditions identified by LTMG-DGE are more significantly enriched to the pathways such as glycolysis, TCA cycle and respiration chain, apoptosis, and lipid metabolism pathways, as well as genes regulated by HIF1A and STAT3 (Figure 5B and Supplementary Table 7). Note that APE1/Ref-1 directly interacts with HIF1A and STAT3 [50, 51], and regulates oxidative stress response, glucose and lipid metabolism, and relevant mitochondrial functions. These results suggest LTMG-DGE method can detect more functionally relevant genes than other tested methods. Complete pathway enrichment results of the differentially expressed genes identified by the tested methods were given in Supplementary Table 7.

We utilized qPCR to investigate 12 selected differentially expressed genes with highest significances identified by LTMG-DGE and MAST, and 7 genes commonly identified by both methods (**Methods**). Specifically, comparing APE1/Ref-1-KD vs. control under hypoxia, (1) nine genes namely STAT3, CREM, SP1, USP3, CDS1, ACTR1A, PARP4, TMEM144, and MNAT1 were identified as down-regulated genes by LTMG-DGE, while not detected as with a significant difference by MAST; (2) three genes namely SEM1, PARPBP and RAP2C were identified as up-regulated by MAST while not with a significant difference by LTMG-DGE; (3) two genes namely MKI67 and TMPO were identified as up-regulated genes by both methods; and (4) five genes namely JUNB, LYPLAL1, PRDM1, PGK1 and SLC2A3 were identified as down-regulated genes by both methods (Figure 5C). Using qPCR, we demonstrated that eight out of the nine genes identified as significantly down-regulated in the scRNA-seq data are confirmed to be down-regulated (p<1e-3 and fold change<0.7), while the three genes identified as up-regulated by MAST but unchanged by LTMG-DGE are truly unchanged in the qPCR experiment. In addition, qPCR confirmed the up- and down-regulation for the seven common differentially expressed genes. These observations clearly suggest a better sensitivity and specificity of LTMG-DGE compared with MAST. To further validate the nine down regulated genes specifically identified by LTMG-DGE, we checked their expression in TCGA pancreatic cancer data and identified eight of the nine are significantly down regulated in the samples with low APE1 expression comparing to the samples with high APE1 expression (p<1e-3 by Mann Whitney test, Supplementary Figure 6).

With the qPCR, we validated 13 down regulated genes namely JUNB, LYPLAL1, PRDM1, PGK1, SLC2A3, STAT3, CREM, USP3, CDS1, ACTR1A, PARP4, TMEM144, and MNAT1. We further checked the differential expression status of these genes identified by SCDE, SC2P, EdgeR and DESeq (all with p<0.01 as the significance cut-off). Down regulation of 0, 11, 0 and 3 out of the 13 genes were identified by SCDE, SC2P, EdgeR and DESeq, respectively.

We also examined the SE and AE peaks for the genes regulated by different TFs. Specifically, in APE1/Ref-1-KD vs. control under hypoxia, LTMG-DGE identified that genes regulated by STAT3 have a higher proportion of SE peaks (Figure 5D-E) while genes regulated by HIF1 have an emerging AE peak with a low-valued mean (Figure 5F). This again implies a regulatory functional loss of STAT3 and attenuation of HIF1 in the APE1/Ref-1-KD cells. Figure 5D-F showcased the gene expression profile and inferred LTMG distribution of PARP4 regulated by STAT3 and SLC2A3 regulated HIF1.

LTMG-GCR was further applied for gene co-regulation analysis. Two co-regulation modules corresponding to the AE of the STAT3 and HIF1A regulated genes and three co-regulation modules corresponding to the SE of the STAT3 and HIF1A regulated genes were identified (Figure 5G and supplementary table 8). Further analysis revealed that the 16 out of the 17 cells of the SE modules are APE1/Ref-1-KD samples and 16 out of the 18 cells of the the AE modules are the control samples, respectively, suggesting a switch of the TRS of STAT3 and HIF1A in the APE1 knock down cells. More interestingly, the AE module of the HIF1A regulated genes include glycolytic genes ALDOA, PGK1 and LDHA, while the two SE modules of HIF1A regulated genes are enriched by genes related to DNA methylation, angiogenesis and other transcriptional factors, which are independent to glycolytic genes, suggesting losing of APE1 results in a suppression of certain HIF1A regulated genes.

We also compared LTMG-GCR with SCENIC [14]. Comparing to LTMG-GCR, Scenic uses the gene co-expression correlation derived from all cells to identify gene co-regulations in scRNA-seq data. In the scenic derived gene coregulation modules, no module regulated by STAT3 was found while only seven genes were identified in the HIF1A regulated module, none of which is related to glycolysis, TCA cycle, or angiogenesis. In addition, majority of down regulated genes in the APE1/Ref-1-KD cells under hypoxia condition were identified in the modules of JUNB and JUND, which we identified as the downstream of STAT3 and HIF1A. Based on our mathematical consideration, we believe the local low rank formulation utilized in LTMG-GCR can better characterize the genes and cells sharing a common transcriptional regulatory signal in the whole gene expression profile, which are determined by of the varied transcriptional regulatory inputs through cells.

## DISCUSSION

We developed LTMG as a statistical model that specifically fits the distribution of scRNA-Seq data. LTMG considers the heterogeneity of transcriptional regulatory states, metabolism rates of mRNA molecules, and experimental resolution in modeling scRNA-seq data. Our comprehensive model evaluations demonstrated that LTMG can accurately infer the multi-modality of genes expression, better handle low expressions caused by suppressed regulation and incompletely degraded mRNA, and has a significantly improved goodness of fitting, compared to other existing models. Our experimental validation suggested the differential gene expression tests LTMG-DGE has better sensitivity and specificity compared to five state-of-art methods. In addition, LTMG-DGE is equipped with a generalized linear model that could deal with comparisons under complex experimental design.

LTMG is designed for analysis of scRNA-seq with a comparable sequencing depth for each cell, and the application of LTMG on drop-seq based data such as 10x Genomics data also demonstrated the model out performs other models in goodness of fitting and can successfully infer multimodality from single gene’s expression profile. However, since there is always a wide span of total reads among the cells in the drop-seq data, in which case, the distribution of the normalized gene expression may be severely affected by variations in total sequenced reads. SC2P introduced a concept to model scRNA-seq considering a cell wise sequencing resolution [52]. A possible future direction of LTMG is to implement a similar cell wise factor into the current LTMG model, so it will improve the characterization of varied expression resolution for drop-seq based scRNA-Seq data. In addition, the inference varied expression state relies on sample size. For the cells collected from a pure condition, on average, LTMG only identified 200-1500 genes with more than one distinct AE peaks when the sample size is several hundreds, while more than 2000 of such genes can be identified when the sample size is larger than 5000.

ScRNA-Seq provides an ideal environment for studying the transcriptional regulatory mechanism, as each gene’s expression in a single cell is the end product of all its current transcriptional regulatory inputs. A key challenge here is to identify the data patterns encoded in scRNA-seq data that corresponds to heterogeneous regulatory signals. LTMG delineates the diversity of the gene expression states of each gene and assign each gene’s expression state in each cell by the Gaussian peak with maximal likelihood, which naturally characterize the regulatory states on single gene and single cell level. This serves as an informative starting point for characterization of gene co-regulation modules. And indeed, application of LTMG-GCR on the APEX1 data demonstrated that modules displaying a bi-clustering structure can be effectively identified and achieved higher specificity comparing to Scenic in a scRNA-seq data set with transcriptional perturbation. The bi-clustering formulation identifies a submatrix in which each gene is with a consistent expression state, which indicates the genes possible co-regulated by a same transcriptional signal specific to a subset of samples, i.e. a local rank-1 submatrix in the complete matrix. We consider the inference of gene co-regulation module via de novo identification of local low rank submatrix is more rational than using the co-expression dependency through all cells, since each gene regulation signal is specific to a certain but unknown subset of cells.

Comparing to the local rank-1 co-regulation module formulation, there are more complicated scenarios. For single cells collected from a highly dynamic biological process, such as cells under fast differentiation, a continuous switch of transcriptional regulatory signals such as phase transitions and delayed effects may result in more complicated expression patterns, which forms a local low rank submatrix instead of a local rank-1 matrix of simple “ON” and “OFF” switch. We anticipate that our LTMG model and its future synthesis with sophisticated low rank structure detection methods, will effectively identify co-regulation modules that stand out in complicated expression patterns caused by incessant switches of all transcriptional regulation states.

Our analysis also suggested that the dimension reduction and cell clustering analysis conducted on LTMG inferred gene expression states better characterize the difference among cell types. Our explanation is that the cell type specifically expressed genes trend to form distinct gene expression states while the general cell physiological state related genes such as cell proliferation and metabolism genes form one peaks of large variance. Hence the LTMG based gene expressions states transformation can identify the genes with most significant varied states, which are more likely to be cell type specific markers. Actually, regulation of the cell type specific genes are with more constant regulatory inputs, which best fit to the assumption of LTMG model (see Supplementary Methods). Successfully distinguishing the cell type and phenotypic genes not only increase the specificity of cell type clustering analysis, but also helps to explain the low rank space of scRNA-seq data and provide more biological meaningful visualization. With the bi-state property observed from transcriptional bursting, we also derived LTMG model can fit the transcriptional bursting regulations. However, a more detailed derivation is needed for the conditions that the multi-modality inferred by LTMG can achieve high the specificity. A straightforward link between LTMG inferred peaks and the transcriptional bursting model is that the proportion and mean of each peak directly correspond to the frequency and expression level of each input signal [53]. Eventually, we hope the LTMG model based inference of gene expression states will shed new light on deducing the mechanisms transcriptional regulation by using scRNA-seq data.

## ACKNOWLEDGEMENTS

C.Z and C.S specifically thank Dr. Yunlong Liu and Dr. Xiongbin Lu from Indiana University for their advice in this work. C.Z thank Dr. Tao Sheng from the University of Georgia and Dr. Xin Chen from Tianjin University for their help in the early stage of this work. C.Z and M.F thank Dr. Mark Kelley from Indiana University School of Medicine for this advice in this study.

## FUNDING SUPPORTS

NIGMS 1R01GM131399-01; NCI 2R01CA167291-06; Showalter Young Investigator Award.

## REFERENCES

1. Puram, S.V., et al., Single-cell transcriptomic analysis of primary and metastatic tumor ecosystems in head and neck cancer. Cell, 2017. 171(7): p. 1611–1624. e24.

2. Azizi, E., et al., Single-cell map of diverse immune phenotypes in the breast tumor microenvironment. Cell, 2018. 174(5): p. 1293–1308. e36.

3. Zheng, G.X., et al., Massively parallel digital transcriptional profiling of single cells. Nature communications, 2017. 8: p. 14049.

4. Finak, G., et al., MAST: a flexible statistical framework for assessing transcriptional changes and characterizing heterogeneity in single-cell RNA sequencing data. Genome biology, 2015. 16(1): p. 278.

5. Vu, T.N., et al., Beta-Poisson model for single-cell RNA-seq data analyses. Bioinformatics, 2016. 32(14): p. 2128–2135.

6. Li, W.V. and J.J.J.N.c. Li, An accurate and robust imputation method scImpute for single-cell RNA-seq data. 2018. 9(1): p. 997.

7. Anders, S. and W.J.G.b. Huber, Differential expression analysis for sequence count data. 2010. 11(10): p. R106.

8. Wang, T., et al., Comparative analysis of differential gene expression analysis tools for single-cell RNA sequencing data. 2019. 20(1): p. 40.

9. Kharchenko, P.V., L. Silberstein, and D.T. Scadden, Bayesian approach to single-cell differential expression analysis. Nature methods, 2014. 11(7): p. 740.

10. Wu, Z., et al., Two-phase differential expression analysis for single cell RNA-seq. 2018. 34(19): p. 3340–3348.

11. McCarthy, D.J., Y. Chen, and G.K.J.N.a.r. Smyth, Differential expression analysis of multifactor RNA-Seq experiments with respect to biological variation. 2012. 40(10): p. 4288–4297.

12. Love, M.I., W. Huber, and S.J.G.b. Anders, Moderated estimation of fold change and dispersion for RNA-seq data with DESeq2. 2014. 15(12): p. 550.

13. Kiselev, V.Y., et al., SC3: consensus clustering of single-cell RNA-seq data. 2017. 14(5): p. 483.

14. Aibar, S., et al., SCENIC: single-cell regulatory network inference and clustering. 2017. 14(11): p. 1083.

15. Becht, E., et al., Dimensionality reduction for visualizing single-cell data using UMAP. 2019. 37(1): p. 38.

16. Wang, B., et al., Visualization and analysis of single-cell RNA-seq data by kernel-based similarity learning. 2017. 14(4): p. 414.

17. Zhang, Y., et al., QUBIC: a bioconductor package for qualitative biclustering analysis of gene co-expression data. 2016. 33(3): p. 450–452.

18. Xie, J., et al., QUBIC2: A novel biclustering algorithm for large-scale bulk RNA-sequencing and single-cell RNA-sequencing data analysis. 2018.

19. Chen, K.H., et al., Spatially resolved, highly multiplexed RNA profiling in single cells. 2015. 348(6233): p. aaa6090.

20. Torre, E., et al., Rare cell detection by single-cell RNA sequencing as guided by single-molecule RNA FISH. 2018. 6(2): p. 171–179. e5.

21. Shah, S., et al., In situ transcription profiling of single cells reveals spatial organization of cells in the mouse hippocampus. 2016. 92(2): p. 342–357.

22. Maston, G.A., S.K. Evans, and M.R. Green, Transcriptional regulatory elements in the human genome. Annu. Rev. Genomics Hum. Genet., 2006. 7: p. 29–59.

23. Lee, T.I. and R.A.J.C. Young, Transcriptional regulation and its misregulation in disease. 2013. 152(6): p. 1237–1251.

24. Ay, A., D.N.J.C.r.i.b. Arnosti, and m. biology, Mathematical modeling of gene expression: a guide for the perplexed biologist. 2011. 46(2): p. 137–151.

25. Khanin, R., et al., Statistical reconstruction of transcription factor activity using Michaelis–Menten kinetics. 2007. 63(3): p. 816–823.

26. Duren, Z., et al., Modeling gene regulation from paired expression and chromatin accessibility data. 2017. 114(25): p. E4914–E4923.

27. van Hijum, S.A., M.H. Medema, and O.P.J.M.M.B.R. Kuipers, Mechanisms and evolution of control logic in prokaryotic transcriptional regulation. 2009. 73(3): p. 481–509.

28. Dar, R.D., et al., Transcriptional burst frequency and burst size are equally modulated across the human genome. 2012. 109(43): p. 17454–17459.

29. Schwanhäusser, B., et al., Global quantification of mammalian gene expression control. Nature, 2011. 473(7347): p. 337.

30. Vanlandewijck, M., et al., A molecular atlas of cell types and zonation in the brain vasculature. 2018. 554(7693): p. 475.

31. He, L., et al., Single-cell RNA sequencing of mouse brain and lung vascular and vessel-associated cell types. Scientific data, 2018. 5: p. 180160.

32. He, L., et al., Single-cell RNA sequencing of mouse brain and lung vascular and vessel-associated cell types. 2018. 5: p. 180160.

33. Dong, J., et al., Single-cell RNA-seq analysis unveils a prevalent epithelial/mesenchymal hybrid state during mouse organogenesis. 2018. 19(1): p. 31.

34. Butler, A., et al., Integrating single-cell transcriptomic data across different conditions, technologies, and species. 2018. 36(5): p. 411.

35. Racle, J., et al., Simultaneous enumeration of cancer and immune cell types from bulk tumor gene expression data. Elife, 2017. 6: p. e26476.

36. Puram, S.V., et al., Single-cell transcriptomic analysis of primary and metastatic tumor ecosystems in head and neck cancer. 2017. 171(7): p. 1611–1624. e24.

37. Jones, S., et al., Core signaling pathways in human pancreatic cancers revealed by global genomic analyses. 2008.

38. Fishel, M.L., et al., Apurinic/apyrimidinic endonuclease/redox factor-1 (APE1/Ref-1) redox function negatively regulates NRF2. 2015. 290(5): p. 3057–3068.

39. Li, G., et al., QUBIC: a qualitative biclustering algorithm for analyses of gene expression data. 2009. 37(15): p. e101–e101.

40. Liberzon, A., et al., Molecular signatures database (MSigDB) 3.0. 2011. 27(12): p. 1739–1740.

41. Wang, J., W.W. Tsang, and G. Marsaglia, Evaluating Kolmogorov’s distribution. Journal of Statistical Software, 2003. 8(18).

42. Zheng, C., et al., Landscape of infiltrating T cells in liver cancer revealed by single-cell sequencing. 2017. 169(7): p. 1342–1356. e16.

43. Zhang, L., et al., Lineage tracking reveals dynamic relationships of T cells in colorectal cancer. 2018. 564(7735): p. 268.

44. Guo, X., et al., Global characterization of T cells in non-small-cell lung cancer by single-cell sequencing. 2018. 24(7): p. 978.

45. Dong, J., et al., Single-cell RNA-seq analysis unveils a prevalent epithelial/mesenchymal hybrid state during mouse organogenesis. Genome biology, 2018. 19(1): p. 31.

46. Barry, M. and R.C. Bleackley, Cytotoxic T lymphocytes: all roads lead to death. Nature Reviews Immunology, 2002. 2(6): p. 401.

47. Guo, Y., et al., Granzyme K degrades the redox/DNA repair enzyme Ape1 to trigger oxidative stress of target cells leading to cytotoxicity. Molecular immunology, 2008. 45(8): p. 2225–2235.

48. Wherry, E.J., T cell exhaustion. Nature immunology, 2011. 12(6): p. 492.

49. Kelley, M.R., M.M. Georgiadis, and M.L. Fishel, APE1/Ref-1 role in redox signaling: translational applications of targeting the redox function of the DNA repair/redox protein APE1/Ref-1. Curr Mol Pharmacol, 2012. 5(1): p. 36–53.

50. Shah, F., et al., APE1/Ref-1 knockdown in pancreatic ductal adenocarcinoma– characterizing gene expression changes and identifying novel pathways using single-cell RNA sequencing. 2017. 11(12): p. 1711–1732.

51. Logsdon, D.P., et al., Regulation of HIF1α under Hypoxia by APE1/Ref-1 Impacts CA9 Expression: Dual-Targeting in Patient-Derived 3D Pancreatic Cancer Models. 2016: p. molcanther. 0253.2016.

52. Wu, Z., et al., Two-phase differential expression analysis for single cell RNA-seq. Bioinformatics, 2018. 1: p. 9.

53. Larsson, A.J., et al., Genomic encoding of transcriptional burst kinetics. 2019. 565(7738): p. 251.

