## Supplementary material for "LTMG: A novel statistical modeling of transcriptional expression states in single-cell RNA-Seq data": Supp File

**SUPPLEMENTARY NOTES**

**Mathematical modeling of gene expression in a single cell from a perspective of transcriptional regulation**

A gene’s expression in a mammalian cell is regulated by the interactions between its DNA molecule and a collection of transcriptional regulatory inputs (TRIs) including (1) transcriptional regulatory factors (TFs) (cis-regulation), (2) miRNA or lncRNA, (3) enhancer and super-enhancer, and (4) epigenetic regulatory signals such histone modification, chromatin folding and DNA methylation. For a gene with *P* possible transcriptional regulation inputs $\mathrm{TRI}_{i}, i=1,\ldots,P$, the probability of its promoter being bound by an RNA polymerase $P_{b}$, which is proportional to the transcriptional rate, can be modeled by a Michaelis Menten model^11^:

$$P_{b}=\frac{R_{0}+\frac{R_{1}\left[ \mathrm{TRI}_{1} \right]}{K_{1}}+\ldots\frac{R_{N}\left[ \mathrm{TRI}_{P} \right]}{K_{N}}+\frac{R_{1,2}\left[ \mathrm{TRI}_{1} \right]\left[ \mathrm{TRI}_{2} \right]}{K_{1,2}}+\ldots+\frac{R_{1,\ldots,N}\left[ \mathrm{TRI}_{1} \right]\left[ \mathrm{TRI}_{2} \right]\ldots\left[ \mathrm{TRI}_{P} \right]}{K_{1,2,\ldots,P}}}{1+\frac{\left[ \mathrm{TRI}_{1} \right]}{K_{1}}+\ldots\frac{\left[ \mathrm{TRI}_{P} \right]}{K_{N}}+\frac{\left[ \mathrm{TRI}_{1} \right]\left[ \mathrm{TRI}_{2} \right]}{K_{1,2}}+\ldots+\frac{\left[ \mathrm{TRI}_{1} \right]\left[ \mathrm{TRI}_{2} \right]\ldots\left[ \mathrm{TRI}_{P} \right]}{K_{1,2,\ldots,P}}}=\frac{\sum_{\Omega\in M\left\{ 1\ldots P \right\}} \frac{R_{\Omega}}{K_{\Omega}}\prod_{i\in\Omega} \left[ \mathrm{TRI}_{i} \right]}{\sum_{\Omega\in M\left\{ 1\ldots P \right\}} \frac{1}{K_{\Omega}}\prod_{i\in\Omega} \left[ \mathrm{TRI}_{i} \right]}, (1)$$

where $R_{i}, \left[ \mathrm{TRI}_{i} \right], K_{i}$ denote production rate, concentration and kinetic parameters associated with the *i*th TRI;$M\left\{ 1\ldots P \right\}$ is the power set of $\left\{ 1\ldots P \right\}$, $\Omega$ denotes an element in $M\left\{ 1\ldots P \right\}$; $R_{\Omega}, K_{\Omega}$denote the production rate and kinetic parameters associated with the subset of TRIs in $\Omega$. Specifically, we call each $\Omega$ as a transcriptional regulatory state (TRS), which is determined by the combination of its TRIs, and reflected by the observed expression in a single cell. Noting that in a single cell the state of each TRI can be rationally simplified to either bound ON or OFF to the DNA molecule, thus the $\mathrm{TRI}_{i}$ is a Boolean variable and the equation (1) is a step function with at most $|M\left\{ 1\ldots P \right\}|=2^{P}$ plateau levels:

$$P_{b}\left( \mathrm{Current} \mathrm{TRS}=\left\{ \mathrm{TRI}_{i}, i\in\Omega\right\} \right)=P_{b}\left( \left\{ \left[ \mathrm{TRI}_{i} \right]\gg0,[\mathrm{TRI}_{j}]=0| i\in\mathcal{M}, j\notin\Omega\right\} \right)=R_{\Omega} (2)$$

For a mammalian cell, the total number of possible TRS can be substantially large due to the diverse number and types of TRS, especially the epigenetic regulators. With this derivation, the number of a gene’s regulatory states is at most $2^{p}$, where p is the number of TRSs of the gene. However, the number of TRSs of a gene in a single cell RNA-seq experiment is always much smaller, majorly due to (i) the phenotypic diversity of the cells measured in one experiment is relatively small, (ii) master repressors such as chromatin folding or certain TFs that can totally suppress the gene’s expression, (iii) multiple suppressive regulation mechanisms will be considered as one due to they are less distinguishable with observed expression level.

Here we specifically discuss that with introducing a Gaussian error to the formula (2), a mixture Gaussian distribution can accurately characterize the varied gene expression states from observed gene expression, via deriving the relationship between varied transcriptional regulatory inputs, transcription rate, mRNA abundance, and observed gene expression signal in each cell.

Denote the expression level of a gene X as $G^{X}(t)$, the probability of X’s promoter being bound by an RNA polymerase at time $t$ as $P_{b}^{X}(t)$, i.e. its transcriptional rate at time t, and its mRNA degradation rate as $D^{X}\left( t \right)$. Noting that the transcriptional rate is proportional to the $P_{b}$ in a single cell, and assuming $D^{X}\left( t \right)\propto G^{X}\left( t \right)$, we have

$$G^{X}\left( t \right)\propto\int\left( P_{b}^{X}\left( t \right)-C^{X}G^{X}\left( t \right) \right)\mathrm{dt} (3)$$

, where $C^{X}$ is the kinetic parameters of the mRNA degradation of X in the current cell, and by equation (2), we have $P_{b}^{X}\left( t \right)=\left\{ R_{\Omega_{i}}^{X} \right|i=1\ldots K corresponds to each TRS\}$, in which ${R_{\Omega}}_{i}$is the RNA polymerase binding probability corresponds to the TRS i. Based on (3), X’s expression level through time interval $[T_{0}, T_{N}]$ can be written as:

$$G_{[T_{0},T_{N}]}^{X}\triangleq\int_{T_{0}}^{T_{N}} \left( P_{b}^{X}\left( t \right)-C^{X}G^{X}\left( t \right) \right)\mathrm{dt}=\sum_{k=0,\ldots,N-1} \int_{T_{k}}^{T_{k+1}} \left( R_{{\Omega_{i}}_{k}}^{X}-C^{X}G^{X}\left( t \right) \right)\mathrm{dt}$$

, in which the gene’s regulatory input at $\left[ T_{k},T_{k+1} \right]$ is $R_{{\Omega_{i}}_{k}}^{X}$.

(i) **Constant regulation input:** The first common understanding of transcriptional regulation is a constant input condition, i.e. each transcriptional regulatory input last long enough to ensure a steady state, under which the gene expression level is $G^{X}=\frac{R_{{\Omega_{i}}_{k}}^{X}}{C^{X}}$ when the transcription rate is $R_{{\Omega_{i}}_{k}}^{X}$. Under this condition, the observations made through $[T_{0},T_{N}$], other than the one in the transition between the k and k+1 regulatory input, are sampled from K constants, where K is the number of different values in $R_{{\Omega_{i}}_{k}}^{X}, k=1\ldots N$. These observations follow a mixture Gaussian distribution of K components after introducing a Gaussian error. For the observations sampled in the transition between the k and k+1 regulatory input signal, we quired the transcription rate, mRNA half-life, and protein half-life. Specifically, the transcription rate, mRNA half-life and protein half-life are at 1-5mins, 100mins, and 1000mins level (1-3), suggesting the turnover rate of transcriptional regulation is much faster than mRNA and protein level changes. Hence the ratio of such observation can be assumed small. In addition, due to $\int_{T_{k}}^{T_{k+1}} \left( R_{{\Omega_{i}}_{k}}^{X}-C^{X}G^{X}\left( t \right) \right)\mathrm{dt}$ is an exponential function of t, the log transformation of the sampling made between two steady states: $\frac{R_{{\Omega_{i}}_{k}}^{X}}{C^{X}}$ and $\frac{R_{{\Omega_{i}}_{k+1}}^{X}}{C^{X}}$are uniformly sampled in $[\min\left( \frac{R_{{\Omega_{i}}_{k}}^{X}}{C^{X}}, \frac{R_{{\Omega_{i}}_{k+1}}^{X}}{C^{X}} \right),\max\left( \frac{R_{{\Omega_{i}}_{k}}^{X}}{C^{X}}, \frac{R_{{\Omega_{i}}_{k+1}}^{X}}{C^{X}} \right)]$ that cannot lead to an extra Gaussian peak if the sampling rate is low. Some transcriptional regulatory signal can have faster turnover than transcriptional factor such as randomness of chromatin motion. For such mechanisms, the ones with high randomness can be assumed as extra errors that modeled in the Gaussian error and the functional ones are discussed in (iii).
(ii) **Transcriptional bursting model:** Recent studies revealed a significant number of genes in bacterial and mammalian are regulated through transcriptional bursting, in which the transcriptional regulatory signals occur as “bursts” in a short term to perturb the gene’s expression. Several recent studies modeled the transcriptional bursting into bi-state a model that the gene expression observed through consecutive transcriptional burst regulations switch between on/off states (4). Considering the different on states of different regulatory signals, current models of the transcriptional burst is consistent with the equation (2), hence can be covered by a mixture Gaussian model. Actually, under the “bursting” condition, the production of mRNA in a short interval is $R_{{\Omega_{i}}_{k}}^{X}\cdot t_{i_{k}}$, where $t_{i_{k}}$ and $R_{{\Omega_{i}}_{k}}^{X}$ are the length and turnover of the $i_{k}$ burst. The time interval between two transcriptional bursts can be varied through several to hundreds mins scale (5) while each burst may last several mins. Considering the mRNA half-life is on 100mins levels, which is much larger than the last of a burst, hence the gene expression observed after a burst event can be considered as sampling through the mRNA production to the end of mRNA’s degradation, the distribution of which is a uniform distribution on the log-transformed mRNA abundance. Under this circumstance, since the uniform distribution does not lead to an additional peak, hence a peak inferred by a mixture Gaussian distribution still indicate an expression state with high specificity.
(iii) **Highly dynamic regulatory signals:** We also consider some genes can be with highly dynamic regulatory signals and fast turnover rate. For these genes, no specific state can be modeled due to the highly perturbed input signals. We consider no expression states can be effectively inferred from this type of gene. However, in the LTMG scheme, the expression part of this gene will still be modeled as a left truncated Gaussian peak with a large variance or a mixture of a SE and an AE peak due to the bi-state property of gene expression.

Denote ${\overset{̃}{x}}_{j}, j=1\ldots N$ as the normalized gene expression level (such as CPM or TPM) of gene X in a scRNA-seq experiment with individual library constructed for N cells and measured with a large sequencing amount. With the above considerations of mRNA dynamics and experimental resolution, we can derive a natural relationship between the repertoire of the TRSs of a single gene and its gene expression profile observed in a scRNA-seq experiment for both of the Constant regulation input and Transcriptional Bursting models (as shown in Figure 1). Specifically, (i) ${\overset{̃}{x}}_{j}$ can unbiasedly reflect the $G_{\Omega_{i}}^{X}$ of the active expression states of X in cell j with a log-Gaussian error when ${\overset{̃}{x}}_{j}$ is large; (ii) an observed low (non-zero) level of ${\overset{̃}{x}}_{j}$ can be caused by multiple reasons including true zero expression with a sequencing error, suppressed regulation of X, and un-fully degraded mRNA after the switch from an active expression state to a suppressed expression state in the cell j; and (iii) the observed zero value of ${\overset{̃}{x}}_{j}$ can be true zero expression or a suppressed expression state undetected by the experiment resolution. To overcome the undistinguishable TRS and dynamics errors of low values of ${\overset{̃}{x}}_{j}$, we introduce a latent threshold ${Zcut}^{X}$ depending on both suppressed expression state and degradation of X. When $\log({\overset{̃}{x}}_{j})>{Zcut}^{X}$, ${\overset{̃}{x}}_{j}$ can reflect $G_{\Omega_{i}}^{X}$ with a log-Gaussian error, while the multimodality of X cannot be reliably inferred when l$\mathrm{og}({\overset{̃}{x}}_{j})\leq{Zcut}^{X}$. Figure 1 shows the relationship between the expression state of a gene, $\log({\overset{̃}{x}}_{j})$, and the threshold ${Zcut}^{X}$. For the $\log({\overset{̃}{x}}_{j})>{Zcut}^{X}$, the values follow a mixture Gaussian distribution truncated by the left of ${Zcut}^{X}$, with each Gaussian peak corresponds to one AE peak. For the l$\mathrm{og}({\overset{̃}{x}}_{j})\leq{Zcut}^{X}$, the values could be from one or multiple SE peaks or true zeros.

It is noteworthy that even the multimodality among SE peaks and true zeros cannot be accurately identified when the sample size is not large enough, one Gaussian peak can effectively characterize the boundary between SE and AE peaks. Specifically, considering a gene with K active expression (AE) and K’ suppressed expression (SE) states through N cells, a scRNA-seq experiment has the resolution to detect the gene expression level of K AEs. However, sometimes the experimental resolution cannot effectively detect the rest K’ SE states since the distribution of the SE states over ${Zcut}^{X}$ is small and are not distinguishable. Hence in total, the LTMG may model the gene expression profile as a mixed Gaussian distribution with K+1 peaks. The rationale of such a consideration is that the outcome of all the TRS with mean expression value lower than ${Zcut}^{X}$ cannot be effective observed with the resolution of the scRNA-seq data set, and be distinguished from 0 observations, hence are all considered as one SE peak with a large variance.

Specifically, we classified the observed gene expressions and its underlying gene expression states into six groups, as shown in Figure 1B. Table 1 details the definition of each class. The peak on the most left side is regarded as a Suppressed Expression peak if its mean is lower than ${Zcut}^{X}$. Each of the Gaussian peaks with mean larger than ${Zcut}^{X}$ is considered as one AE peak. With such a definition, one gene can have 0-M AE peaks and 0 or 1 SE peak, where M is pre-defined maximal number of possible gene expressions states of a gene.

Table 1. Definition of the six regions defined in Figure 1,

| Observed gene expression\ Gene expression states | Suppressed Expression State | Activated Expression State |
| --- | --- | --- |
| Below $Z_{\mathrm{cut}}^{X}$ | (1) True Non-Expression of the SE (multiple SE signals are considered as one SE peak in the model) | (2) Undetected True Expressions from the AE |
| Above $Z_{\mathrm{cut}}^{X}$, below the boundary between the SE and most left AE peak | (3) SE state & un-fully degraded mRNA above the experimental resolution | (4) Type II error of True expression of the AE |
| Above the boundary between the SE state and most left SE peaks | (5) Type I error of SE state & un-fully degraded mRNA | (6) True AE expression |

**Benchmarks from previous studies**

Our derivation suggests that the identifiable AE states of X form a truncated mixture distribution on the right of ${Zcut}^{X}$. Multiple established models have confirmed the necessity of using a multi-modal distribution to characterize single gene’s expression profile in scRNA-seq data. Table 2 listed the commonly used methods and models for scRNA-seq gene expression modeling and differential gene expression tests. In detail, SCDE uses a fixed Poisson distribution to model the drop out events and a negative binomial distribution to model the expression part of a gene. MAST considers zero observations as dropouts and the rest as following a log-normal distribution. Trung Nghia Vu et al considers the underlying transcriptional heterogeneity can be fitted by a Beta distribution and utilized a zero-inflated Beta-Poisson distribution for gene expression modeling. SC2P assumes all the genes in one cell share the same zero-inflated negative binomial distribution for the dropout events ^10^. It is noteworthy that both SCDE and SC2P aims to derive a common parameter set to model the dropouts for all the genes in one or multiple cell samples; MAST only considers the zero parts in (1) and (4) in Figure 1 as dropouts; and Beta-Poisson considers the centers of $G^{X}$ through the cells follow a Beta distribution. However, none of the models links the observed expression profile with the underlying regulators of the gene expression—heterogeneity of gene expression states and mRNA metabolism.

Based on our derivation from a perspective of the dynamics of transcriptional regulation and gene expression states, as shown in Figure 1, expressions in part (1) and (4) are unreliable for inference of the distributions, part (2) and (6) are the gene expression of SE states or un-fully degraded mRNAs, the boundary of which depends on the transcriptional dynamics and mRNA degradation of the gene, and (3) and (5) are from an AE state of the gene. Hence, theoretically, it is necessary to have a gene-wise consideration of $Q^{X}$, SE and AE states for accurate modeling of individual gene expression profile in scRNA-seq data.

To validate the LTMG model and its downstream analysis, we compared the model with existing ones on 23 high-quality scRNA-seq data sets collected from the public domain. We also generated a scRNA-seq data with qPCR experiment to experimentally validate the modeling hypothesis and LTMG based differential gene expression test. Our key observations include (1) LTMG achieves a significant better goodness of fitting comparing to the Beta-Poisson, Zero-Inflated Gaussian (MAST) and Zero-Inflated Mixture Gaussian models in more than 90% of the analyzed data sets, (2) LTMG more accurately characterizes cell type specifically expressed genes compared to other models, (3) LTMG based differential gene expression analysis has a better identification accuracy than the existing methods. In addition, the system biology assumption of the LTMG model is validated by significant associations between the fitted LTMG parameters with experimentally measured mRNA kinetics. The full computational approach including LTMG fitting and downstream analysis are released through an open source R package.

Existing methods include SCDE, MAST, Beta-Poisson, scImpute, and SC2P focusing on optimizing the best parametric form of the “dropout” proportions in the data. Although all these models admitted the necessity of a multimodal assumption, none of the methods links the observed gene expression level with their underlying regulators – the heterogeneous transcriptional regulatory states and mRNA transcription/degradation kinetics. Specifically, a more flexible assumption of the multimodality is needed to handle the genes regulated by multiple transcriptional regulators.

Table 2. Comparison of existing models

|  | **#parameters** | **Consideration of Dropout** | **Consideration of expression** | **Data** |
| --- | --- | --- | --- | --- |
| **SCDE** | 3 | Fixed Poisson | Negative Binomial | Counts |
| **MAST** | 3 | Zero inflation | Log Gaussian | TPM/CPM/… |
| **Beta-Poisson** | 4 | 0 inflated beta-Poisson | beta-Poisson | TPM/CPM/… |
| **SC2P** | 5 | Sample wise Negative Binomial | Log Gaussian | Counts |
| **ScImpute** | 5 | Log Gamma | Log Gaussian | TPM/CPM/… |
| **LTMG-2LR** | 5 | gene wise censored data | Log Gaussian | TPM/CPM/… |
| **LTMG** | Depends on fitting | gene wise censored data | Log Mixture Gaussian | TPM/CPM/… |

With the LTMG model and its link with gene expression states, we define *a gene is* *differentially expressed between the cells of two conditions, if at least one* gene expression state *(either SE or AE) of the gene is with a significant different representing level in one condition versus the other*. With this definition, we formulate the computation of a differentially expressed gene (DEGs) in a scRNA-seq data by two steps – (1) identifying all the gene expression states of the gene through the cells of the to-be-tested conditions, and (2) a statistical test of if one gene expression state is with a significantly higher representation rate in one of the to-be-tested conditions.

Our comprehensive analysis revealed that on average more than 83.8% genes in the PC and CC groups are fitted with one and two peaks, which can be well fitted by an LTMG-2LR model. In order to (1) guarantee the mathematical rigorousness, (2) maximize the statistical power and (3) ensure the implementation of a GLM model for a multiple or crossed condition test, we developed a statistical testing approach by using the LTMG-2LR model for the genes with one or two fitted gene expression states, and the LTMG model for the genes with more than two expression states (see detailed derivations in Methods). For a given gene X in a scRNA-seq data of J conditions, denote

$$X_{j}=\left\{ x_{i}^{j}, i=1\ldots N_{j} \right\}, j=1\ldots J$$

as its expression profile in the $N_{j}$ cells of the jth conditions. The following pseudo codes illustrate our differential gene expression analysis approach, namely LTMG-DEG, with detailed formulations of each step given in Method.

**A discussion of the distribution used to cover AE and SE expression parts and evaluation methods**

The distinct feature of LTMG is to model multimodality of the high expression part with considering the low expression below the experimental resolution as left censored data. The left truncation assumption can well handle the complicated errors of the zero and low expression part to avoid over fittings in multimodality inference. As discussed above, existing models such as SCDE, MAST, Beta-Poisson, scImpute, and SC2P have different assumptions for the zero and low expression. To benchmark LTMG and compare with these methods, two factors namely goodness of fitting and model complexity need to be evaluated.

We consider there are multiple ways to compare distributions, such KS statistics, BIC, and likelihood ratio test. Each method is with certain advantage and drawback with respect to the models that are going to be compared for. Likelihood ratio test is rigorous; however, it tests if a growth of model complexity may make a significant improvement to the likelihood, hence is not suitable to compare models of different assumptions. Both likelihood ratio test and BIC can evaluate model complexity while KS only evaluates goodness of fitting, which can suffer an over fitting problem. BIC has been applied to compare different models, however, for the models of single cell RNA-seq, BIC has certain limitations as detailed below:

(a) All zero inflated and left truncated models are with an unbiased estimation of the probability (density) for zero observations, however, Beta-Poisson and Gamma Normal models underestimate the probability of zero observations due to the continuous assumption, which inflate the likelihood of non-zero observations. Zero inflated models consider zero as a discrete distribution and left truncated model consider zero as censored data, hence a BIC based comparison between left truncated and zero-inflated model can be only made on the non-zero part. However, when compare with BP and GN models, the inflated likelihood of non-zero observations may cause a biased evaluation. The inflation of the pdf of non-zero observations can be seen on KS plot of the non-zero expression part.
(b) The effect of model complexity can be reduced when the comparison of KS was made on the models with a same number of parameters, such as comparing Gamma-Normal, two peak LTMG and Beta-Poisson with five parameters. Hence the comparison of KS of ZIG vs LTMG and ZIMG models should also consider model complexity.
(c) We consider the most straightforward way to compare the distributions is through a visualization of fitted pdf and the KS plot. We provided visualization of the plots by the Github link: <https://github.com/zy26/LTMGSCA/tree/master/compare>
(d) BIC is used in the selection of best peak number for the LTMG and ZIMG model.

Based on these considerations, we considered two rational comparisons, include (1) using BIC to compare LTMG and other zero inflated models, and (2) using KS statistics to compare LTMG and other non-zero inflated models only on the genes fitted with the same number of parameters in each case, such that the models being compared have the same complexity level. For (1), we compared the BIC of LTMG, ZIMG and ZIG (Mast model) on selected testing data set. No surprisingly, LTMG achieved the best BIC comparing to these two zero inflated models, due to ZIMG may consider the zero as one component and non-zero low expression as one or multiple peak that has a higher model complexity than LTMG while ZIG cannot effectively handle the genes multimodality AE states. For (2), we compared the LTMG with Beta Poisson (five parameters version) and Gamma Gaussian model used in scImpute, both of which are with five parameters, only on the genes fitted with five parameters by LTMG, majority of which are with one SE and one AE peak. We identified the LTMG consistently achieved better KS statistics comparing to the two other models. Detailed visualization of the overall comparison and the pdf of each distribution fitted to each gene are given in the Github link: <https://github.com/zy26/LTMGSCA/tree/master/compare>

**SUPPLEMENTARY METHODS**

**Datasets processing and classification details**

We processed and classified cells into different groups, Complete Data (CD), Cell Cluster (CC) and Pure Condition (PC) for the purpose of evaluating LTMG and other models in a comprehensive manner. Complete Data (CD) is considered as a mixture of different cells. It could be cells from a patient, cells of a tissue or just all the cells within a dataset. Cell Cluster (CC) is considered as a group of functionally similar cells clustered by gene expression value. For instance, in a head and neck cancer dataset (GSE103322), the authors differentiated sequenced cells into malignant, T cells, endothelial cells, etc. Furthermore, as indicated by the original papers, malignant cells are highly heterogenetic between different patients. In that case, malignant cells alone could be divided by original patients. For a fair comparison, we did not include any cell cluster that was clustered by us, all the cell cluster information is directly retrieved from the original paper. Intuitively different datasets applied different cluster algorithms with different parameters, in order to cope with the complexity of different tissues or samples. It would be very reckless if we just utilize a simple algorithm and the same set of parameters to cluster cells. Meanwhile, the performance comparison of models on already clustered cells could reflect the robustness of models as these clusters coming from different models and parameters. Pure Condition (PC) is regarded as a group of cells in a pure biological setting, such as cells from the same cell line, cells sorted by surface markers. Altogether we collected 20 human single cell RNA seq datasets, and further generated testing data sets by splitting cells within each dataset into CD, CC and PC groups. Detailed processing and classification methods are as followed:

*GSE114397:* This dataset sequenced breast cancer cell line that undergoes EMT. Cells as a whole are regards as CD.

*GSE89567:* This dataset is about IDH-mutant astrocytoma. Cells as a whole are regarded as CD.

*GSE104276:* This dataset is about human prefrontal cortex in different developmental stages. Such that cells are a mixture of different cells. We regarded this dataset as CD.

*GSE110686:* This dataset sequenced CD3+ T cell. Cells as a whole are regarded as CD.

*GSE89497:* This dataset is about human placenta. Cells as a whole are regarded as CD.

*10X_PBMC:* This dataset sequenced peripheral blood cell. Cells as a whole are regarded CD.

*GSE76312:* This dataset is about stem cells in chronic myeloid leukemia. Cells are also classified by different patient mutation (positive or negative). Classified cells are still a mixture, they are regarded as CD.

*GSE98638:* This dataset is about T cells in cancer. Cells can be sorted by the origin of the patient. They further clustered cells in several sub clusters. Cells classified by the patient are regarded as CD. Cells clustered by expression are regarded as CC.

*GSE70630:* This dataset is about oligodendroglia. Cells from different patients are considered as CD.

*GSE84789:* This dataset is about B cells in NSCLC. B cells are classified as from normal sample and tumor sample, denoted as NB and TB. Sorted by FACS, NB and TB are considered as PC.

*GS75688:* This dataset is about human primary breast cancer. We classified cells by different patient mutation type in breast type. These classified cells are still a mixture of different cells so that they are regarded as CD.

*GSE89232:* This dataset is about human pre-cDCs. Cells as a whole are regarded as CD. Cells sorted by surface marker are regarded as PC.

*GSE102130:* This dataset is about cells from K27M-mutant glioma patient. We did not find specific cell type annotation information. But cells could be classified based on the origin of different patients. Cells classified by patients are considered as CD.

*GSE94820:* This dataset is about human blood dendritic cells, monocytes and progenitors. Cells (CD141, double negative, CD1C, and pDC) are sorted by surface marker, thus are regarded as PC. Cells are later classified by gene expression using a clustering method like AXLSIGLEC6, C100C34int are regarded as CC.

*GSE103322:* This dataset is about head and neck cancer. Cell type and patient information are provided in the original paper. Cells classified by cell type are regards as CC. Cells classified from patients are considered as CD.

*GSE100618:* This dataset is about human hemopoietic lymphoid-myeloid progenitor, including LMPP, MLP, GMP. Cells are sorted by surface marker, using flow cytometry. Classified LMPP, MLP and GMP cells are considered as PC.

*GSE72056:* This dataset is about human melanoma. Cells from different patients are considered as CD. Authors of the original paper also provided cell type information, clustered with gene expression data. These classified cells are regarded as CC.

*GSE84465:* This dataset is about glioblastoma. Cells as a whole are considered as CD. Cells clustered by gene expression are considered as CC.

*GSE99305:* This dataset is about Pancreatic Ductal Adenocarcinoma with a focus on APE1. Cells as a whole are considered as CD.

*GSE81861:* This dataset is about human colorectal tumors. Cells sorted by the origin of patients are considered as CD. Cells from specific cell line are considered as PC.

*GSE86618:* This dataset is about epithelial cells in Idiopathic Pulmonary Fibrosis. Cells are sorted by FACS using an epithelial marker (positive), hematopoietic marker (negative) and endothelial marker (negative). Sorted cells can be also classified from the origin of patients. Thus, we regarded sorted cells from the same patient as PC.

*GSE70580:* This dataset is about human tonsil innate lymphoid cells (ILCs). Cells are sorted by the surface marker. Patient information is also provided. Cells in each patient are considered as CD, but sorted cells in each patient are considered as PC.

*GSE106540:* This dataset is about the transformation of T cells under different simulations. This dataset as a whole is regarded as CD. Cells sorted by surface marker are considered as PC.

Altogether we collected 23 human datasets dated from January 2016 to June 2018 with a sample size between 100 and 7500 cells. Majorities of them are from 2000 to 6000 cells. To be noted, after classification, some cluster or pure condition may be removed due to low sample size, like myocyte cells in dataset GSE103322 (only 19 cells). Moreover, we also set a 200 genes cutoff for each classified dataset to make sure our comparison is comprehensive and just. That is if we applied three or four models on a dataset, but we could only get less than 200 genes with valid KS value for all the models. We will remove that dataset in the comparison analysis. After all, we get 51 PC, 49 CC and 78 CD datasets in our comparison.

**EM algorithm to estimate the parameters of the LTMG model**

Denoting the log-transformed RPKM expression level of a gene X over N cells as $X=\left( x_{1}, x_{2}, \ldots,x_{N} \right)$. We assume that $x_{i}$ follows a mixture Gaussian distribution with K Gaussian peaks corresponding to different SE and AE peaks. We introduce a parameter $Z_{\mathrm{cut}}^{X}$ and consider the log transformed zero and low expression values smaller than $Z_{\mathrm{cut}}$ as left censored data. With the left truncation assumption,$X$ is divided into reliably measured expressions ($x_{j}\geq Z_{\mathrm{cut}}^{X}$) and left-censored gene expressions ($x_{j}<Z_{\mathrm{cut}}^{X}$). The likelihood function of X can be written as:

$$p\left( X | \Theta\right)=\prod_{j=1}^{N} p\left( x_{j} | \Theta\right)=\prod_{j=1}^{M} \sum_{i=1}^{K} a_{i}p_{i}\left( x_{j} | \theta_{i},x_{j}\geq Z_{\mathrm{cut}}^{X} \right)\cdot\prod_{j=M+1}^{N} \sum_{i=1}^{K} a_{i}p_{i}\left( x_{j} | \theta_{i},x_{j}<Z_{\mathrm{cut}}^{X} \right)=\prod_{j=1}^{M} \sum_{i=1}^{K} a_{i}\frac{1}{\sqrt{2\pi\sigma_{i}}}e^{-\frac{\left( x_{j}-\mu_{i} \right)^{2}}{2\sigma_{i}^{2}}}\cdot\prod_{j=M+1}^{N} \sum_{i=1}^{K} a_{i}p_{i}\left( x_{j} | \theta_{i},x_{j}<Z_{\mathrm{cut}}^{X} \right)=L\left( \Theta| X \right) (*)$$

, where parameters $\Theta=\left\{ a_{i}, u_{i} \sigma_{i} \right| i=1\ldots K\}$ and $a_{i}, u_{i} and \sigma_{i}$ are the mixing probability, mean and standard deviation of the K Gaussian distributions, corresponding to K expression states, M is the number of observations $x_{j}$ that are larger than $Z_{\mathrm{cut}}^{X}$, N is the total number of observations. $\Theta$ can be estimated using EM algorithm with given $Z_{\mathrm{cut}}^{X}$ and K. An EM algorithm to estimate the parameters $\Theta=(\alpha_{i}, u_{i}, \sigma_{i}| i=1\ldots K)$ of an LTMG model treats latent variables $y_{i} \mathrm{and} Z_{j}$ as missing data. With pre-specified K and $Z_{\mathrm{cut}}^{X}$, the Q function is defined as follows:

$$\mathcal{Q}\left( \Theta^{t},\Theta^{t-1} \right)=\sum_{i=1}^{K} \sum_{j=1}^{M} \log(\alpha_{i}^{t}p_{i}\left( x_{j} | u_{i}^{t},\sigma_{i}^{t} \right))p(y_{j}=i|x_{j},\Theta^{t-1})+$$

$$\sum_{j=M+1}^{N} \int_{-\infty}^{Z_{\mathrm{cut}}^{X}} \sum_{i=1}^{K} \log(\alpha_{i}^{t}p_{i}(Z_{j}|u_{i}^{t},\sigma_{i}^{t}))p({y_{j}=i|Z}_{j},\Theta^{t-1})p(Z_{j}|x_{j},\Theta^{t-1})dZ_{j}$$

, where $p_{i}\left( x_{j} | u_{i}^{t},\sigma_{i}^{t} \right)=\frac{1}{\sqrt{2\pi\sigma_{i}^{t}}}e^{-\frac{\left( x_{j}-u_{i}^{t} \right)^{2}}{2{\sigma_{i}^{t}}^{2}}}$, $Z_{j}$is a latent variable for the true value of $x_{j}$ if $x_{j}<Z_{\mathrm{cut}}^{X}$, i.e. left censored data, $y_{j}=1\ldots K$ are latent variables indicating that $x_{j}$ comes from the $j$th Gaussian peak, and *t* is the current iteration step, M is the number of observations $x_{j}$ that are larger than $Z_{\mathrm{cut}}^{X}$, N is the total number of observations. The Q function considers a mixture Gaussian likelihood function for the $x_{j}\geq Z_{\mathrm{cut}}^{X}$ and $Z_{j}$, the latent true value of $x_{j}$ when $x_{j}<Z_{\mathrm{cut}}^{X}$, as described in the equation (*). And $Z_{j}$ follows a distribution of the left truncated part of the mixture Gaussian distribution. For the $t\mathrm{th}$ iteration of the algorithm, with computed parameter set $\Theta^{t-1}$, the pdf of $Z_{j}$ and probability of $y_{j}$are:

$$f\left( Z_{j}|\Theta^{t-1} \right)=\frac{\sum_{i=1\ldots K} \frac{1}{\sqrt{2\pi\sigma_{i}^{t-1}}}e^{-\frac{\left( Z_{j}-u_{i}^{t-1} \right)^{2}}{2{\sigma_{i}^{t-1}}^{2}}}}{\sum_{i=1\ldots K} \int_{-\infty}^{Z_{\mathrm{cut}}^{X}} \frac{1}{\sqrt{2\pi\sigma_{i}^{t-1}}}e^{-\frac{\left( y-u_{i}^{t-1} \right)^{2}}{2{\sigma_{i}^{t-1}}^{2}}}dy}, Z_{j}\in(-\infty,Z_{\mathrm{cut}}^{X})$$

$$p\left( y_{j}=K|\Theta^{t-1},x_{j}\geq Z_{\mathrm{cut}}^{X} \right)=\frac{\frac{1}{\sqrt{2\pi\sigma_{k}^{t-1}}}e^{-\frac{\left( x_{j}-u_{k}^{t-1} \right)^{2}}{2{\sigma_{k}^{t-1}}^{2}}}}{\sum_{i=1\ldots K} \frac{1}{\sqrt{2\pi\sigma_{i}^{t-1}}}e^{-\frac{\left( x_{j}-u_{i}^{t-1} \right)^{2}}{2{\sigma_{i}^{t-1}}^{2}}}}, k=1\ldots K$$

$$p\left( y_{j}=k|\Theta^{t-1},x_{j}<Z_{\mathrm{cut}}^{X} \right)=\frac{\int_{-\infty}^{Z_{\mathrm{cut}}^{X}} \frac{1}{\sqrt{2\pi\sigma_{k}^{t-1}}}e^{-\frac{\left( y-u_{k}^{t-1} \right)^{2}}{2{\sigma_{k}^{t-1}}^{2}}}dy}{\sum_{i=1\ldots K} \int_{-\infty}^{Z_{\mathrm{cut}}^{X}} \frac{1}{\sqrt{2\pi\sigma_{i}^{t-1}}}e^{-\frac{\left( y-u_{i}^{t-1} \right)^{2}}{2{\sigma_{i}^{t-1}}^{2}}}dy}, k=1\ldots K$$

The parameter $\Theta$ are estimated by the following EM algorithm:

$$\begin{matrix} Q\left( \Theta,\Theta^{t-1} \right)=\sum_{i=1}^{K} \sum_{j=1}^{M} \log\left( a_{i}p_{i}\left( x_{j}|\mu_{i},\sigma_{i} \right) \right)p\left( y_{j}=i|x_{j},\Theta^{t-1} \right) \\ +\sum_{j=M+1}^{N} \int\sum_{i=1}^{K} \log\left( a_{i}p_{i}\left( Z_{j}|\mu_{i},\sigma_{i} \right) \right)p\left( y_{j}=i|Z_{j},\Theta^{t-1} \right)p\left( Z_{j}|x_{j},\Theta^{t-1} \right)dZ_{j} \\ =\sum_{i=1}^{K} \sum_{j=1}^{M} \log\left( a_{i} \right)p\left( y_{j}=i|x_{j},\Theta^{t-1} \right) \\ +\sum_{i=1}^{K} \sum_{j=1}^{M} \log\left( p_{i}\left( x_{j}|\mu_{i},\sigma_{i} \right) \right)p\left( y_{j}=i|x_{j},\Theta^{t-1} \right) \\ +\sum_{i=1}^{K} \sum_{j=M+1}^{N} \int\log\left( p_{i}\left( x_{j}|\mu_{i},\sigma_{i} \right) \right)p\left( Z_{j}|x_{j},\Theta^{t-1} \right)dZ_{j}p\left( y_{j}=i|x_{j},\Theta^{t-1} \right) \\ =\sum_{i=1}^{K} \sum_{j=1}^{M} \log\left( a_{i} \right)p\left( y_{j}=i|x_{j},\Theta^{t-1} \right) \\ +\sum_{i=1}^{K} \sum_{j=1}^{M} \log\left( p_{i}\left( x_{j}|\mu_{i},\sigma_{i} \right) \right)p\left( y_{j}=i|x_{j},\Theta^{t-1} \right) \\ +\sum_{i=1}^{K} \sum_{j=1}^{N} \log\left( a_{i} \right)p\left( y_{j}=i|Z_{j},\Theta^{t-1} \right) \\ +\sum_{i=1}^{K} \sum_{j=M+1}^{N} \frac{1}{2\sigma_{i}^{2}}p\left( y_{j}=i|x_{j},\Theta^{t-1} \right)\left[ E\left( Z_{j}^{2}|\mu_{i}^{t-1},\sigma_{i}^{t-1},Z_{j}<Z_{cut} \right)-2\mu_{i}E\left( Z_{j}|\mu_{i}^{t-1},\sigma_{i}^{t-1},Z_{j}<Z_{cut} \right)+\mu_{i}^{2} \right], \end{matrix}$$

The M step can be written as:

$$\frac{\partial Q}{\partial a_{i}}=0\to a_{i}^{t}=\frac{1}{N}(\sum_{j=1}^{M} P\left( i \right|x_{j},\Theta^{t-1})+\sum_{j=M+1}^{N} P(i|Z_{j},Z_{cut},\Theta^{t-1}))$$

$$\frac{\partial Q}{\partial\mu_{i}}=0\to\mu_{i}^{t}=\frac{\sum_{j=1}^{M} x_{j}P\left( i|x_{j},\Theta^{t-1} \right)+\sum_{j=M+1}^{N} \left( \mu_{i}^{t-1}-\sigma_{i}^{t-1}H\left( \frac{Z_{cut}-\mu_{i}^{t-1}}{\sigma_{i}^{t-1}} \right) \right)P\left( i|Z_{j},Z_{cut},\Theta^{t-1} \right)}{\sum_{j=1}^{M} P(i|x_{j},\Theta^{t-1})+\sum_{j=M+1}^{N} P(i|Z_{j},Z_{cut},\Theta^{t-1})}$$

$$\frac{\partial Q}{\partial\sigma_{i}}=0\to\sigma_{i}^{t^{2}}=\frac{\sum_{j=1}^{M} P\left( i|x_{j},\Theta^{t-1} \right)\left( x_{j}-\mu_{i}^{t-1} \right)^{2}+\sigma_{i}^{\left( t-1 \right)^{2}}\sum_{j=M+1}^{N} \left( 1-\frac{Z_{cut}-\mu_{i}^{t-1}}{\sigma_{i}^{t-1}}*H\left( \frac{Z_{cut}-\mu_{i}^{t-1}}{\sigma_{i}^{t-1}} \right) \right)P(i|Z_{j},Z_{cut},\Theta^{t-1})}{\sum_{j=1}^{M} P(i|x_{j},\Theta^{t-1})+\sum_{j=M+1}^{N} P(i|Z_{j},Z_{cut},\Theta^{t-1})},$$

where

$$\Theta^{t-1}=\{a_{i}^{t-1},\mu_{i}^{t-1},\sigma_{i}^{t-1}|i=1\ldots K\},$$

$$P\left( i|Z_{j},Z_{cut},\Theta^{t-1} \right)=\frac{\int_{-\infty}^{Q} P\left( i|x_{j},\Theta^{t-1} \right)}{\sum_{i=1}^{K} \int_{-\infty}^{Q} P\left( i|x_{j},\Theta^{t-1} \right)}=\frac{P\left( -\infty<Z_{j}<Z_{cut}|\mu_{i}^{t-1},\sigma_{i}^{t-1} \right)}{\sum_{i=1}^{K} P\left( -\infty<Z_{j}<Z_{cut}|\mu_{i}^{t-1},\sigma_{i}^{t-1} \right)} ,$$

$$H\left( x \right)=\frac{\phi\left( x \right)}{\Phi\left( x \right)} ,$$

$\phi\left( x \right)$ and $\Phi\left( x \right)$ are the pdf and cdf of standard normal distribution,

$$P\left( i|x_{j},\Theta^{t-1} \right)=\prod_{j=1}^{N} \sum_{i=1}^{K} a_{i}^{t-1}\frac{1}{\sqrt{2\pi}\sigma_{i}^{t-1}}e^{-\frac{\left( x_{j}-\mu_{i}^{t-1} \right)^{2}}{2\left( \sigma_{i}^{t-1} \right)^{2}}}.$$

And the E step is:

$$E\left( Z_{j}|\mu_{i}^{t-1},\sigma_{i}^{t-1},Z_{j}<Z_{cut} \right)=\mu_{i}^{t-1}+\sigma_{i}^{t-1}E\left( \varepsilon_{j}|\varepsilon_{j}<\frac{Z_{cut}-\mu_{i}^{t-1}}{\sigma_{i}^{t-1}} \right)=\mu_{i}^{t-1}+\frac{\sigma_{i}^{t-1}\int_{-\infty}^{\frac{Q-\mu_{i}^{t-1}}{\sigma_{i}^{t-1}}} w\phi\left( w \right)\mathrm{dw}}{\Phi\left( \frac{Z_{cut}-\mu_{i}^{t-1}}{\sigma_{i}^{t-1}} \right)}=\mu_{i}^{t-1}+\frac{\sigma_{i}^{t-1}\phi\left( \frac{Z_{cut}-\mu_{i}^{t-1}}{\sigma_{i}^{t-1}} \right)}{\Phi\left( \frac{Z_{cut}-\mu_{i}^{t-1}}{\sigma_{i}^{t-1}} \right)}=\mu_{i}^{t-1}+\sigma_{i}^{t-1}H(\frac{Z_{cut}-\mu_{i}^{t-1}}{\sigma_{i}^{t-1}}).$$

Similarly,

$$E\left( Z_{j}^{2}|\mu_{i}^{t-1},\sigma_{i}^{t-1},Z_{j}<Z_{cut} \right)=\mu_{i}^{\left( t-1 \right)^{2}}+\sigma_{i}^{\left( t-1 \right)^{2}}-\sigma_{i}^{t-1}(Z_{cut}+\mu_{i}^{t-1})H(\frac{Z_{cut}-\mu_{i}^{t-1}}{\sigma_{i}^{t-1}})$$

$\Theta$ can be estimated by iteratively running the E and M step in the above algorithm with given $Q, X \mathrm{and} K$.

**Selection of K**

After the EM algorithm is executed for $K$ = 1, …, 5, the optimal $K$ is selected with the minimal Bayesian Information Criterion (BIC):

$$BIC=-2\ln\left( \Theta^{*} \right)+3Kln(N)$$

, where $N$ is the number of cell samples. Particularly, LTMG-2LR denotes an LTMG distribution with two peaks, and the mean of one peak is smaller than $Z_{cut}$ while the other larger than $Z_{cut}$, i.e. $\Theta=\left\{ a_{i}, u_{i}, \sigma_{i} \right|i=1, 2\}$, $u_{1}<Z_{cut}$ and $u_{1}>Z_{cut}$. A cell is designated to a mixing component or a peak within which it has the largest probability to fall.

**Determination of** $\mathbf{Z}_{\mathbf{cut}}$

Intuitively, $Z_{cut}$ can be determined by using the minimal non-zero observation in each gene’s expression profile. However, for some housekeeping genes or genes with fast mRNA degradation rate, there are only active and zero expression. Hence the minimal non-zero observation of these genes cannot reflect the experimental resolution. To identify the genes with minimal non-zero observation can truly indicate the experimental resolution. We fitted a mixture Gaussian distribution for the minimal non-zero observation of all the expressed genes in each single cell data (see details in Supplementary Methods). Our analysis suggested that a mixture of two peaks were identified in all the tested data. The peak with the smaller mean corresponds to the true lower bound of the experimental resolution, and the other peak corresponds to the lowest expressed AE peaks of the housekeeping genes. With this observation, we developed the following criteria to determine the $Z_{cut}$ of each gene X. Denote $Z_{Cut}^{0}$ as the boundary of the two Gaussian peaks identified in the above way. If the minimal non-zero observation of X is smaller than $Z_{Cut}^{0}$, $Z_{Cut}^{X}=min(x|x\in X,x>0)$, otherwise, $Z_{Cut}^{X}=Z_{Cut}^{0}$.

**Parameter fitting of the LTMG-2LR model for gene expression data of multiple conditions**

To conduct a rigorous statistical testing for the differentially expressed genes through multiple conditions, we define a LTMG-2LR distribution by an LTMG distribution with two peaks, and the mean of one peak is smaller than $Q$ while the mean of the other peak is larger than $Q$, i.e. $\Theta=\left\{ a_{i}, \mu_{i}, \sigma_{i} \right|i=1, 2\}$, $\mu_{1}<Q$ and $\mu_{1}>Q$. Denote $X_{j}=\left\{ x_{i}^{j}, i=1\ldots N_{j} \right\}, j=1\ldots J$ as its expression profile of gene X in the $N_{j}$ cells of J conditions, an LTMG_2LR is fitted for each $X_{j}$ with assuming the same parameter set ${(u}_{0}^{X},\sigma_{0}^{X})$ of the SE peak through all the conditions as shown below:

$$\left\{ \begin{matrix} X_{1}\sim LTMG\_2LR\left( a_{1}^{X}, \mu_{0}^{X}, \mu_{1}^{X}, \sigma_{0}^{X}, \sigma_{1}^{X} \right) \\ X_{2}\sim LTMG\_2LR\left( a_{2}^{X}, \mu_{0}^{X}, \mu_{2}^{X}, \sigma_{0}^{X}, \sigma_{2}^{X} \right) \\ X_{3}\sim LTMG\_2LR\left( a_{3}^{X}, \mu_{0}^{X}, \mu_{3}^{X}, \sigma_{0}^{X}, \sigma_{3}^{X} \right) \\ \ldots\end{matrix} \right..$$

A modified EM algorithm is developed for parameter estimation of the above formulation. Specifically, the parameters are estimated by the following Q function and modified M steps:

$$Q^{*}=\sum_{j=1}^{J} Q\left( \Theta_{j}^{t},\Theta_{j}^{t-1},X_{j} \right), \Theta^{t}=\{a_{i}^{t},\mu_{i}^{t},\sigma_{i}^{t}|i=1\ldots K\}$$

$$\frac{\partial Q^{*}}{\partial a_{j}}=0\to a_{j}^{t}, \mathrm{under} \mathrm{fixed} \{,\sigma_{0}^{t-1},\mu_{j}^{t-1},\sigma_{j}^{t-1}|j=1\ldots J\}$$

$$\frac{\partial Q^{*}}{\partial\mu_{j}}=0\to\mu_{j}^{t}, \mathrm{under} fixed \{\mu_{0}^{t-1},\sigma_{0}^{t-1},\sigma_{j}^{t-1},a_{j}^{t}|j=1\ldots J\}$$

$$\frac{\partial Q^{*}}{\partial\sigma_{j}}=0\to\sigma_{j}^{t}, \mathrm{under} \mathrm{fixed} \{\mu_{0}^{t-1},\sigma_{0}^{t-1},\mu_{j}^{t-1},a_{j}^{t}|j=1\ldots J\}$$

$$\frac{\partial Q^{*}}{\partial\mu_{0}}=0\to\mu_{0}^{t}, \mathrm{under} \mathrm{fixed} \{\sigma_{0}^{t-1},\mu_{j}^{t},\sigma_{j}^{t},a_{j}^{t}|j=1\ldots J\}$$

$$\frac{\partial Q^{*}}{\partial\sigma_{0}}=0\to\sigma_{0}^{t}, \mathrm{under} \mathrm{fixed} \{\mu_{0}^{t-1},\mu_{j}^{t},\sigma_{j}^{t},a_{j}^{t}|j=1\ldots J\}$$

**LTMG based gene co-regulation analysis**

By the formulation of LTMG, for a gene with one SE state and K-1 different AE states, its expression profile across different single cells is modeled by a mixture of K Gaussian distributions; and (2) for a group of genes co-regulated by a specific TRS, each gene’s expression profile, in those cells regulated by the TRS, forms a unimodal Gaussian distribution, after involving independent Gaussian errors. Hence a gene co-regulation model corresponds to a submatrix enriched by 1s in the Binary matrix $M$ constructed in the following way:

For a gene X’s expression profile through N samples fitted with one SE and K-1 AE peaks, denote $P_{i}^{X}=0, 1\ldots K-1, i=1,\ldots,N$as the peak with the highest likelihood

$$L\left( X_{i}, \mathrm{peak} k \right)=a_{k}\frac{1}{\sqrt{2\pi\sigma_{k}}} e^{-\frac{\left( X_{i}-\mu_{k} \right)^{2}}{2\sigma_{k}^{2}}}, i=1\ldots N,$$

in which 0 represents the SE peak and $1\ldots K-1$ represents the AE peaks. Then a ($K-1)\times N$ binary matrix $M_{(K-1)\times N}^{X}$ can be constructed by

$$M_{(K-1)\times N}^{X}\left[ i,j \right]=\left\{ \begin{matrix} 1, \mathrm{if} P_{i}^{X}=j \\ 0, \mathrm{if} P_{i}^{X}\neq j \end{matrix} \right. ,$$

$i=1\ldots N, j=1\ldots K-1$. $M$ is merged by $M^{X}$ for the $X$ with at least one AE peak. A bi-cluster enriched by 1s in $M$ corresponds a group of genes and cells, each of the gene is regulated by one specific TRS through the cells, which is potentially a gene co-regulation module.

We applied our in-house developed bi-clustering method QUBIC with the above binary matrix construction method to build a gene co-regulation module identification method, namely LTMG-GCR. Specifically, the QUBIC is directly implemented from the QUBIC R packaged with the following parameters: -o 3000 -f 0.25 -c 0.95. LTMG-GCR is applied to the scRNA-seq data of APEX/Ref-1 KD experiment. Pathway enrichment analysis of the genes in the identified bi-clusters are computed by using a hypergeometric test against the canonical pathway and transcriptional regulation targets encoded by MsigDB, with p<0.001 as a significant cutoff.

**Model comparisons on uncovering true expression dynamics using single molecule Fish data**

To further validated the multi-modality inferred by LTMG, we collected a data set with scRNA-seq and smFISH data collected from the same cell population (6). To the best of our knowledge, this is the only dataset with smFISH and scRNAseq parallelly measured on the same population of cells under a fixed experimental condition. Due to the limited coverage of smFISH, only 15 genes both identified by smFISH and scRNAseq were utilized in this analysis.

We fitted scRNAseq probability distributions by using three models namely LTMG, ZIMG and MAST(ZIG), and calculated the probability mass function of smFISH data. Noted, both scRNAseq and smFISH data are log normalized. Kullback-Leibler divergence (KL divergence) measures the differences between probability distributions. KL divergence between the PDF of scRNAseq data and the PMF of smFISH data was further computed for each of the 15 genes, by using the following equations:

$$KL(P_{X}\left| Q_{X}^{i} \right)=\sum_{x\in X} P\left( x_{scFISH} \right)\log\left( \frac{P\left( x_{scFISH} \right)}{Q^{i}\left( x_{scRNAseq} \right)} \right)$$

, where $x_{scFISH}$ and $x_{scRNAseq}$ represent the decentralized profile of the smFISH and scRNAseq data of the gene $X$, respectively; $P_{X}$ denotes the PMF of the smFISH profile of gene X; and $Q_{X}^{i}$ denotes the PDF of the scRNAseq profile of X fitted with *i=*LTMG, ZIMG, and MAST.

**Stringent condition for validation of multimodality**

Denote the pdf of the mixture Gaussian distribution inferred from gene expression profile of N cells $X=\left( x_{1}, x_{2}, \ldots,x_{N} \right)$as:

$$pdf\left( x | \Theta\right)=\sum_{i=1}^{K} a_{i}p_{i}\left( x | \theta_{i} \right)=\sum_{i=1}^{K} a_{i}\frac{1}{\sqrt{2\pi\sigma_{i}}}e^{-\frac{\left( x_{j}-\mu_{i} \right)^{2}}{2\sigma_{i}^{2}}}$$

For peak *k*, denote $T_{k}$ as $T_{k}=\{t|\frac{a_{k}\frac{1}{\sqrt{2\pi\sigma_{k}}}e^{-\frac{\left( t-\mu_{k} \right)^{2}}{2\sigma_{k}^{2}}}}{\sum_{i=1}^{K} a_{i}\frac{1}{\sqrt{2\pi\sigma_{i}}}e^{-\frac{\left( t-\mu_{i} \right)^{2}}{2\sigma_{i}^{2}}}}>p_{\mathrm{cutoff}}\}$, i.e. all the real values with a more than $p_{\mathrm{cutoff}}$ probability to belong to the peak *k* by the mixture Gaussian likelihood function. In this study, we use $p_{\mathrm{cutoff}}$=0.95. Then we define the stringent cover of peak *k* as the largest continuous open interval covering $\mu_{k}$in $T_{k}$, i.e. $\mathrm{SC}_{k}=\left( {LB}_{k},{HB}_{k} \right), s.t. \underset{{LB}_{k},{HB}_{k}}{\mathrm{argmax}} ({HB}_{k}-{LB}_{k}),\left( {LB}_{k},{HB}_{k} \right)\in T_{k}, and \mu_{k}\in\left( {LB}_{k},{HB}_{k} \right)$, and $\mathrm{SC}_{k}=\emptyset$ if $\mu_{k}\notin T_{k}$, as illustrated in the figure below, in which the red, green and blue regions are the $\mathrm{SC}_{1}, \mathrm{SC}_{2},$ and $\mathrm{SC}_{3}$ for the peak 1, 2 and 3, and the black region belongs to $T_{2}$ but is excluded in $\mathrm{SC}_{2}$ due to it is not covered by a continuous interval in $T_{2}$ containing $\mu_{2}$.

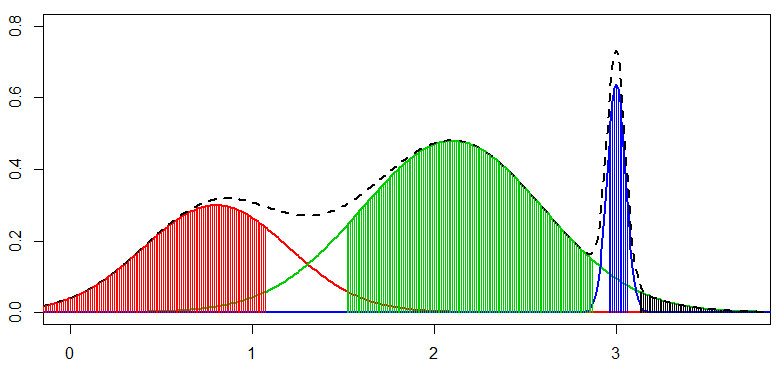

We define peak *k* is distinct to others if there are more than 5% of cells with the $X's$ expression belong to $\mathrm{SC}_{k}$, It is noteworthy that we applied this stringent condition to three large single cell data sets composing more 5000 cells, the 5% of which is with more than 250 cells.

**Measure low dimensional visualization with silhouette width.**

Silhouette width measures the consistency among a group of cells in any embedded low dimension within clusters of data, hence can be utilized to measure the goodness of embedded dimension in representing the cell clusters when the true cell label is given. The silhouette width value represents the overall similarity (distance) of an object to its own cluster compared to other clusters. Detailed mathematic formation of silhouette analysis can be found in [2]. Here we calculated Silhouette width value based their Euclidian distance in the embedded 2D dimension with respect to the known cell labels, by using silhouette function in the cluster R package. We then sum up the Silhouette value of every cell type in a dataset, we denoted sil, as the proxy of the quality of low dimensional visualization in Figure 4.

**LTMG-SCA – an R package for scRNA-seq data analysis within the LTMG scheme**

With an inference of the heterogeneous gene expression states and possible TRS of each gene by our LTMG model, analysis of following problems for a scRNA-seq data, including (1) differentially expressed genes, (2) transcriptional bursts, (3) cell type specific gene regulations, (4) cell type inference, and (5) gene co-regulation module identification, can be formulated and computed by implementing LTMG model with other methods (Table 3 and Supplementary Methods). An R package of LTMG model based single cell RNA-seq analysis tools was developed and released through GitHub: https://github.com/zy26/LTMGSCA. Specifically, the package includes the fitting of LTMG and LTMG-2LR models for single cell RNA-seq data, LTMG-DGE, LTMG-GCR, LTMG based likelihood assessment, outlier gene and cell identification, identification of phenotypic feature associated gene expression states, and cell type inference, as detailed in Table 3.

Table 3. Functions in the package LTMG-SCA

| Question(s) | Biological definition | Formulation within the gene expression states framework | Computational Solution |
| --- | --- | --- | --- |
| Differential gene expression | Genes show differential expression | One gene is with one or several varied AE peaks in a group of cells comparing to others | Implement LTMG with a GLM or enrichment test (LTMG-DEG) |
| Transcription burst identification | High expression caused by a strong transcriptional activation | Expression of a gene that is significantly larger than its largest AE peak | Outlier identification based on LTMG |
| Cell type specific gene regulation | A TRS specific to a cell type | A gene with one or several SE/AE peaks that enrich a certain cell type | correlate the gene expressions states inferred by LTMG with known cell types |
| Cell type inference | A group of cells forms a possible cell type | Cells with a distinct number of cell-type specific gene expression states | Implementation of LTMG with a cell clustering algorithm |
| Gene co-regulation identification | Genes co-regulated by a TRS | A group of genes with consistent SE or AE peaks in a certain subset of cells | Implement LTMG with bi-clustering |

**Supplementary References**

1.	Friedel, C.C., Dölken, L., Ruzsics, Z., Koszinowski, U.H. and Zimmer, R.J.N.a.r. (2009) Conserved principles of mammalian transcriptional regulation revealed by RNA half-life. **37**, e115-e115.

2.	Maiuri, P., Knezevich, A., De Marco, A., Mazza, D., Kula, A., McNally, J.G. and Marcello, A.J.E.r. (2011) Fast transcription rates of RNA polymerase II in human cells. **12**, 1280-1285.

3.	Boisvert, F.-M., Ahmad, Y., Gierliński, M., Charrière, F., Lamont, D., Scott, M., Barton, G., Lamond, A.I.J.M. and Proteomics, C. (2012) A quantitative spatial proteomics analysis of proteome turnover in human cells. **11**, M111. 011429.

4.	Wang, Y., Ni, T., Wang, W. and Liu, F.J.B.R. (2019) Gene transcription in bursting: a unified mode for realizing accuracy and stochasticity. **94**, 248-258.

5.	Corrigan, A.M., Tunnacliffe, E., Cannon, D. and Chubb, J.R.J.E. (2016) A continuum model of transcriptional bursting. **5**, e13051.

6.	Torre, E., Dueck, H., Shaffer, S., Gospocic, J., Gupte, R., Bonasio, R., Kim, J., Murray, J. and Raj, A.J.C.s. (2018) Rare cell detection by single-cell RNA sequencing as guided by single-molecule RNA FISH. **6**, 171-179. e175.

**SUPPLEMENTARY FIGURES AND TABLES**

**Supplementary figure 1.** Heatmap and t-SNE plot of GSE99235 and GSE98816. To identify mouse fibroblast cells, we applied Seurat on two datasets GSE99235 and GSE98816 using default parameters, the detailed heatmap and t-SNE plot are shown above.

**Supplementary figure 2.** KS test on more datasets of different groups. We compared models on a comprehensive dataset. The top 30 datasets in each group are shown in Figure 2. The rest are shown in above.

**Supplementary figure 3.** Linking scRNAseq with smFISH with different models. A) We applied KL divergence to measure the similarity of each model fitted scRNAseq data with smFISH. B) Some example gene distribution on smFISH and scRNAseq.

**Supplementary figure 4.** Fitted peak distribution of liver, lung and colon cancer infiltrated T cells.

**Supplementary figure 5.** Low dimensional visualization of cells with different origin by LTMG t-SNE in head and neck cancer microenvironment.

**Supplementary figure 6.** LTMG-DGE identified down regulated gene by APEX low expression are also down regulated in TCGA pancreatic cancer datasets.

**Supplementary table 1.** Dataset meta information in model comparison analysis.

**Supplementary table 2.** Fitting comparison of multiple datasets (4 models).

**Supplementary table 3.** qPCR results in figure 5C

**Supplementary table 4.** KS value of example datasets in figure 2B.

**Supplementary table 5.** Detailed fitting comparison of genes of different peaks (3 models).

**Supplementary table 6.** Averaged uncensored region proportion.

**Supplementary table 7.** Enrichment comparison of AE, Exp and SE peak.

**Supplementary table 8.** LTMG-DGE, MAST, SCDE, SC2P, EdgeR and DEseq identified differentially expressed genes and enriched pathway.

**Supplementary table 9.** Gene co-regulation modules of STAT3 and HIF1A regulated genes in the APEX1 data.

**Abbreviation List**

scRNA-seq: single-cell RNA-sequencing

LTMG: Left truncated mixture Gaussian distribution

smFISH: single molecular in situ hybridization

SE: suppressed expression

AE: active expression

TRI: transcriptional regulatory input

TRS: transcriptional regulatory state

GES: gene expression state

ZIMG: zero inflated mixture Gaussian distribution

BPSC: Beta-Poisson single-cell distribution

KS: Kolmogorov Smirnov Statistic

GLM: generalized linear model

PDF: probability density function

HNSC: Head-Neck Squamous Cell Carcinoma

LTMG-DGE: LTMG based differential gene expression analysis

LTMG_GCR: LTMG based gene co-regulation analysis

APE1/Ref-1-KD: APE1/Ref-1knockdown

TF: transcriptional regulatory factor
